# A protective role for the cerebellum in cognitive aging

**DOI:** 10.1101/2024.10.15.618102

**Authors:** Federico d’Oleire Uquillas, Esra Sefik, Jakob Seidlitz, Jewel Merriman, Veronica Zhang, Jonathan D. Cohen, Rafael Romero-Garcia, Varun Warrier, Richard A.I. Bethlehem, Aaron F. Alexander-Bloch, Jorge Sepulcre, Samuel S.-H. Wang, Patrizia Vannini, Jesse Gomez

## Abstract

**Background:** Brain reserve — the brain’s resilience to age-related change or damage — provides protection against cognitive decline. The cerebellum is relatively unstudied as a contributor to brain reserve. This study investigates cerebellar brain reserve in the largest cohort to date.

**Methods:** We used data from the Human Connectome Project (n=708, 36-100yrs), UK Biobank (n=45,013, 44-81yrs), and ADNI (n=1,423, 56-95yrs). ADNI participants were cognitively normal or had a diagnosis of mild cognitive impairment or Alzheimer’s disease (AD) dementia. We examined associations between cerebellar tissue volume, age, Montreal Cognitive Assessment (MoCA) scores, global PET amyloid burden, and APOE genotype.

**Findings:** HCP-Aging data revealed heterogenous aging-associated changes in cerebellar volume, with the greatest effects in posterior hemispheric regions (crus I) (Bonferroni-corrected, *p*<0·05). MoCA scores were associated with higher tissue density in the cerebellum (*p*<0·0001) to the same extent as neocortex, and MoCA scores coupled most strongly with posterior cerebellar cortex. Strikingly, greater volume in MoCA visuospatial-related cerebellar cortex protected against aging-related cognitive decline (*p*=0·0001). We replicated tissue aging results in the UK Biobank with the greatest aging-related effects in posterior cerebellum (*p*<0·0001), and an association of greater cerebellar volumes with less cognitive decline (Trails Making-B: *p*<0·00001; Digit Symbol Substitution: *p*=0·034). AD patients with low amyloid-beta burden (Aβ−) exhibited the strongest cerebellar association with MoCA (volume x group, Aβ− AD: *p*=0·0001). In Aβ− individuals, APOE ε4/ε4 carriers showed the greatest effect with MoCA (volume x APOE, ε4/ε4: *p*=0·017).

**Interpretation:** Our large-scale study demonstrates a potentially strong role for the cerebellum in mitigating cognitive decline. The persistence of this protection in APOE ε4/ε4 carriers reshapes our understanding of reserve and AD risk. Our findings open the cerebellum as a novel target for future clinical research on brain reserve in aging populations.

**Funding:** National Science Foundation, National Academies of Sciences, Engineering and Medicine, National Institutes of Health.

**Research in Context:** *Evidence before this study:* We searched PubMed and GoogleScholar between August 12, 2023, and September 25, 2024 for articles irrespective of language or date of publication, relating to measures of cortical and cerebellar aging. Search terms included “cerebellum”, “cognitive aging”, “cerebellar reserve”, “Alzheimer’s disease”, “covariance”, “retrogenesis”, and “connectivity”. Prior research demonstrated that the cerebellum may play a role in cognitive function. Studies have shown that cerebellar development is spatially heterogenous, with posterior regions undergoing the most protracted change in structure and function. Brain reserve has been predominantly studied in the neocortex, with some studies suggesting the cerebellum may contribute to cognitive changes in diseases like Alzheimer’s disease (AD) dementia. Cerebellar volume differences between young and older adults have been noted, as well as between healthy adults and neurodegenerative conditions like Parkinson’s disease. However, the cerebellum’s contribution to cognitive reserve, particularly in healthy aging and individuals with a clinical diagnosis of mild cognitive impairment or AD dementia, had not been explored especially in large population-based datasets.

*Added value of this study:* To our knowledge, this study is the largest to date looking into how cerebellar structures contribute to cognitive outcomes in both healthy older adults, and those with mild cognitive impairment or AD dementia. We leverage three large neuroimaging datasets — HCP-Aging, UK Biobank, and ADNI — and demonstrate that cerebellar aging is spatially heterogeneous, with posterior regions showing the greatest age-related decline. We further establish a significant link between larger cerebellar volumes and better cognitive outcomes, suggesting that the cerebellum plays a role in cognitive resilience. Additionally, we identify how cerebellar structures predict cognitive performance and interact with amyloid-beta brain pathology and APOE genotype, particularly in those with low amyloid brain burden and those at greatest risk of AD such as in homozygous APOE e4 allele carriers.

*Implications of all the available evidence:* The aging of the global population raises the challenge of maintaining cognitive health in older age. Our research focuses on the cerebellum as a novel mediator of preserved cognitive function in old age and clinical dementia. Our findings have profound implications, given that individuals with robust cerebellar structures may be missed during cognitive and clinical screenings. Greater cerebellum volume seems to be most advantageous in those at higher risk for Alzheimer’s disease based on APOE4 status and those who have yet to accumulate substantial amyloid-beta brain pathology. Our study underscores the importance of the cerebellum as a novel brain reserve mechanism in aging populations.

## Introduction

The global demographic shift toward an aging population creates a need to better understand cognitive aging. The number of individuals older than 70 years old is growing more quickly than that of younger adults (Garmany et al., 2021). While the cerebellum has been widely recognized as mediating sensory processing and motor adaptation, it has become recently appreciated for its association with a variety of cognitive domains (King et al., 2019; Adamaszek et al., 2017; Levisohn et al., 2000). Liu et al. (2022) demonstrated that cerebellar cortex develops along a rostrocaudal gradient, with posterior regions contributing to cognitive functions and showing protracted functional and structural maturation. This developmental trajectory mirrors the maturation of cerebral cortex and age-related behavioral improvements (Gaiser et al., 2024), suggesting the cerebellum may contribute to cognitive resilience in later life.

Brain reserve, or the brain’s ability to withstand damage without exhibiting overt clinical symptoms, has traditionally been studied in neocortex (Stern et al., 2019). However, the dysmetria hypothesis (Schmahmann et al., 1998) suggests that cerebellar impairment disrupts cognitive processing, and an aging cerebellum may contribute to cognitive and emotional difficulties in Alzheimer’s disease (AD) (Jacobs et al., 2018). Conversely, evidence suggests that the cerebellum may confer protection (Mitoma et al., 2020). Cerebellar volume differs in younger and older adults (Bernard & Seidler, 2013), and decreased cerebellar volume has been associated with compromised Montreal Cognitive Assessment (MoCA) scores in various neurodegenerative diseases (Kerestes et al., 2023; Ozturk et al., 2021). However, the larger role the cerebellum may play in aging remains untested, and its potential as a source of cognitive reserve holds impact for individuals along the trajectory of mild cognitive impairment or AD.

Leveraging data from the Human Connectome Project (HCP-Aging), the UK Biobank, and the Alzheimer’s Disease Neuroimaging Initiative (ADNI), we examined associations between cerebellar volume, cognition, amyloid-beta plaques, and APOE genotype. By integrating findings across health and disease, we find strong evidence for the cerebellum’s contribution to brain reserve. We focused on three questions: 1) Does the cerebellum show spatial heterogeneity as it ages? 2) Do differences in cerebellar aging relate to individual differences in cognitive scores? 3) How do these processes manifest in clinical conditions like AD dementia?

## Methods

In this cross-sectional study, we sourced MRI data from three major cohorts to investigate cerebellar-cognitive relationships across the adult lifespan: the Human Connectome Project-Aging (HCP-Aging), the UK Biobank, and the Alzheimer’s Disease Neuroimaging Initiative (ADNI). We included a total of 47,144 participants spanning a broad age range. The HCP-Aging cohort included 708 participants aged 36–100 years. The UK Biobank dataset provided 45,013 participants aged 44–81 years, and the ADNI cohort included 1,423 participants aged 56–95 years old (**Table 1**). All participants provided informed consent, and each cohort adhered to institutional review board guidelines.

**Table 1a.**
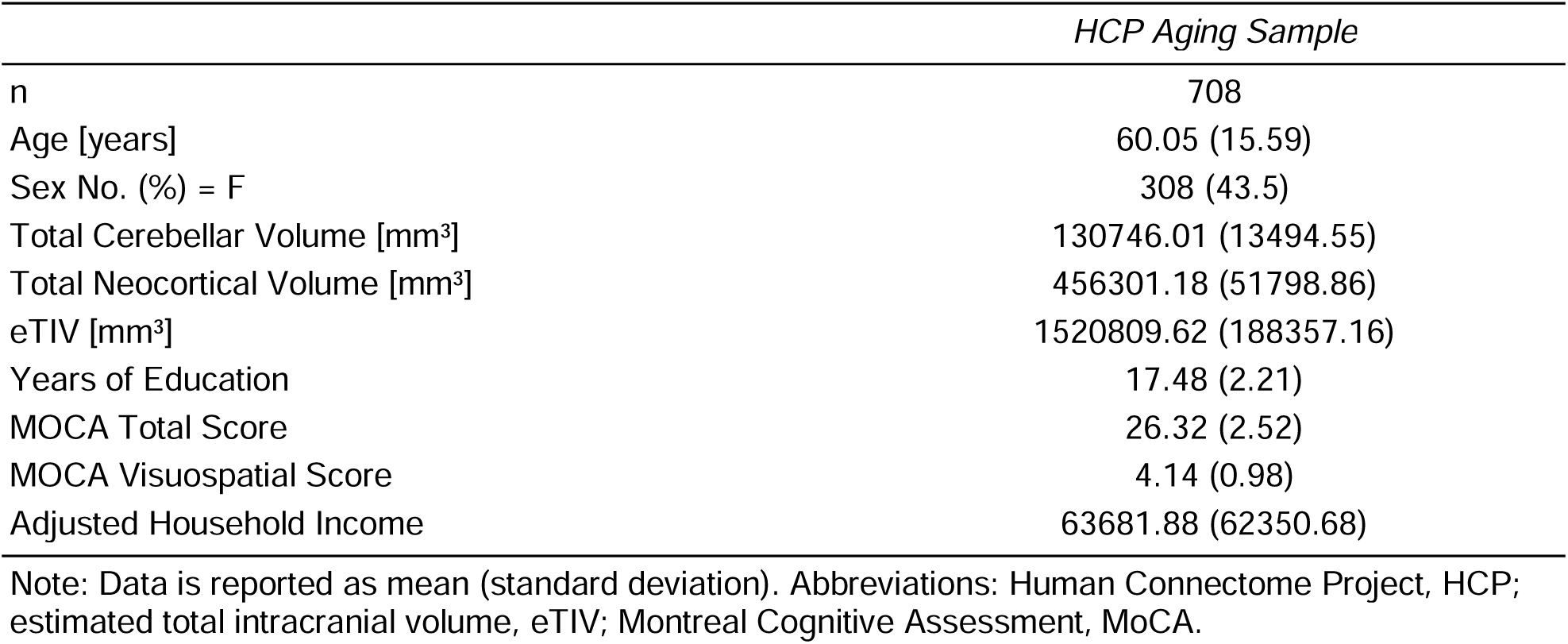
Sample Demographics and Imaging Characteristics in the HCP Aging Sample.

**Table 1b.**
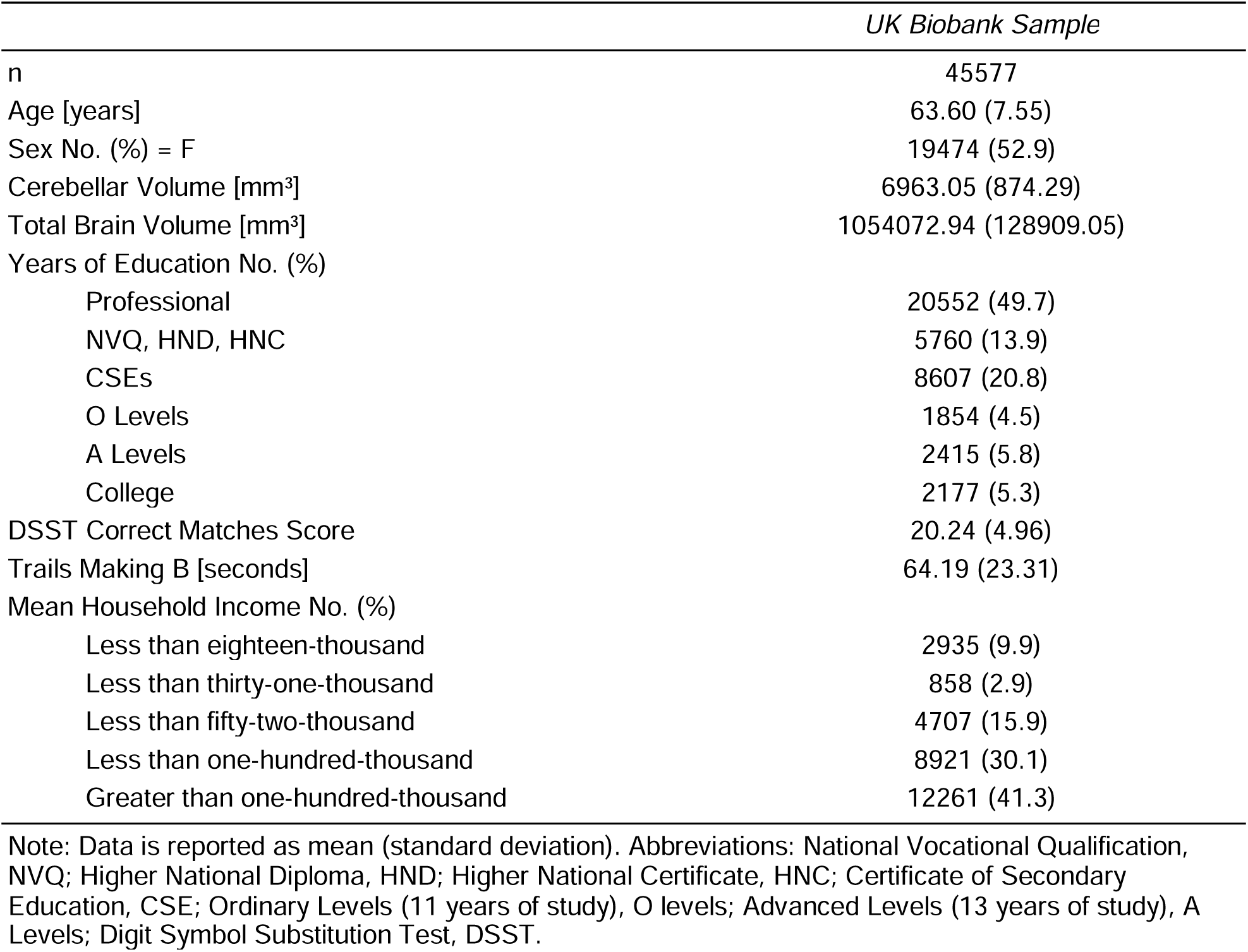
Sample Demographics and Imaging Characteristics in the UK Biobank Sample.

**Table 1c.**
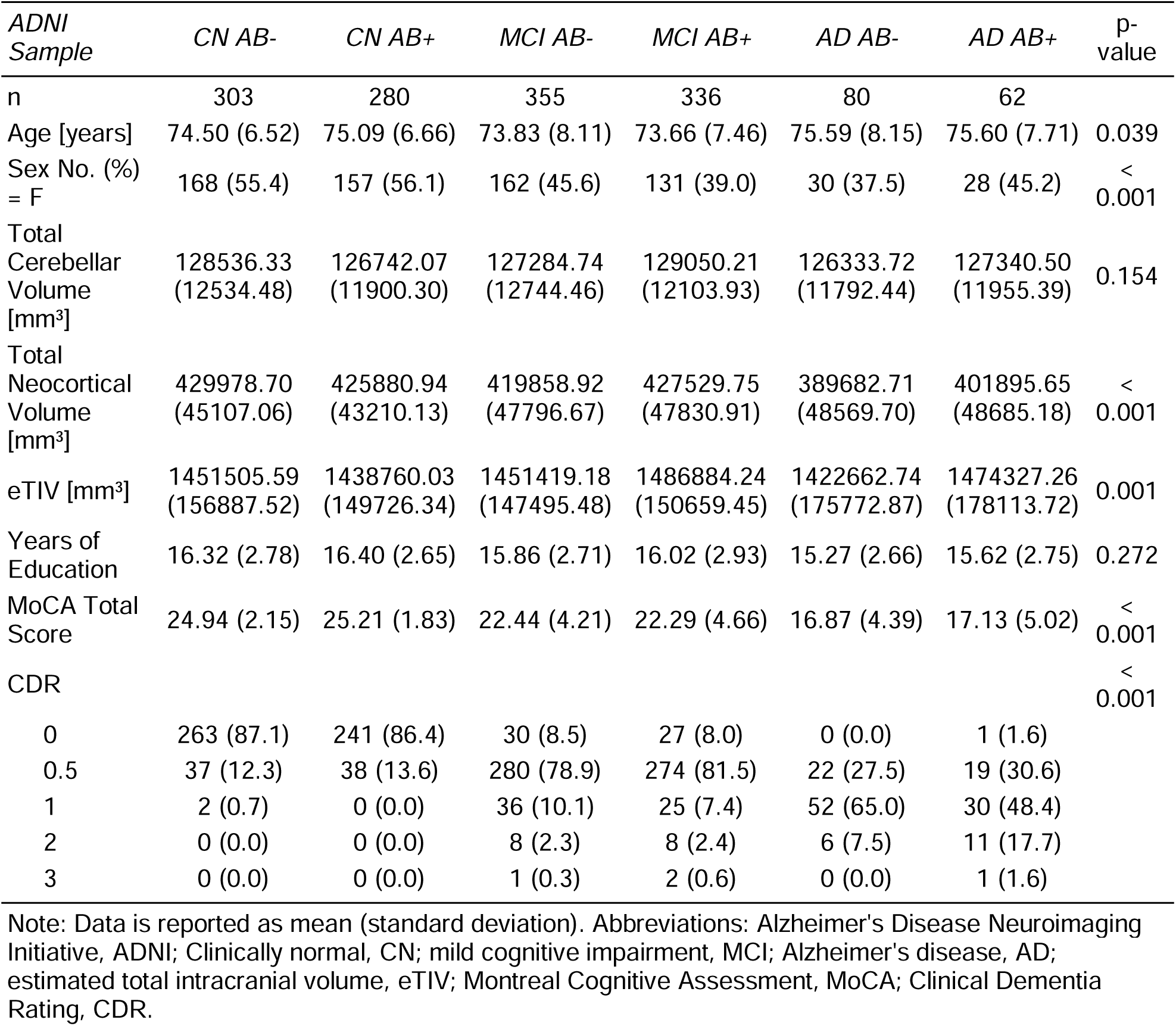
Sample Demographics and Imaging Characteristics in the ADNI Sample.

We included a racially diverse sample of participants with complete cerebellar coverage on MRI scans, as well as corresponding cognitive testing. In the HCP-Aging cohort, there were 2 (0.28%) American Indian/Alaskan Native individuals, 51 (7.2%) Asian, 98 (13.84%) Black or African American, 512 (72.32%) White, 31 (4.38%) more than one race, and 14 (1.98%) individuals with unknown or unreported race, with 74 (10.45%) identifying as Hispanic or Latino, 632 (89.27%) not Hispanic or Latino, and 2 (0.28%) of unreported ethnic group. We also sourced MRI data from the UK Biobank, but did not have access to ethnic or racial status. Additionally, our ADNI cohort comprised of 2 (0.14%) American Indian or Alaskan Native individuals, 7 (0.50%) Asian, 22 (1.55%) Black or African American, 1 (0.07%) Native Hawaiian or other Pacific Islander, 670 (47.08%) White, 11 (0.77%) individuals of more than one race, and 710 (49.89%) individuals of unknown or unreported racial status, with 16 (1.12%) identifying as Hispanic or Latino, 694 (48.77%) not Hispanic or Latino, and 713 (50.11%) of unreported ethnic group.

Cognitive assessments were tailored based on the cohort, and included the Montreal Cognitive Assessment (MoCA) (Nasreddine et al., 2005) within 365 days of MRI procedures, the Digit Symbol Substitution Test (DSST, Wechsler, 1955) and Trails Making B test (TMB, Reitan, 1958), tests of processing speed and attention, and the Clinical Dementia Rating (CDR) scale. Clinical diagnoses in ADNI were determined by a consensus of trained clinicians using cognitive scores and the CDR scale; further details can be found in the Supplemental section. The ADNI cohort encompassed clinically normal (CN), mild cognitive impairment (MCI), and individuals with an Alzheimer’s disease (AD) dementia diagnosis.

All participants underwent T1-weighted MRI scans on 3T MRI scanners. MRI scans were segmented into regions of interest using FreeSurfer (7·1·1) and the specialized ACAPULCO pipeline (Han et al., 2020) for cerebellar cortices for HCP and ADNI cohorts, and FSL for the UK Biobank; detailed image acquisition, processing, and quality control procedures for the UK Biobank can be found in Miller et al. (2016) and Alfaro-Almagro et al. (2018). ADNI also incorporated amyloid-PET with [^11^C]-Pittsburgh compound B lasting 20 minutes (four 5-minute frames) at 50-70 minutes after bolus injection. All PET images were processed and aligned by the ADNI group using FreeSurfer 6·4, and regional uptake value ratios (SUVRs) were calculated from the 50–70-minute post-injection time window, with partial volume correction (PVC). APOE allele genotype in ADNI was determined by blood sample using the KASP genotyping assay.

We quantified associations between sample characteristics using the statistical software package R (v·4·0·4: https://www.R-project.org) (**Table 1**). Multivariate linear regression models were conducted across the HCP, UK Biobank, and ADNI cohorts, covarying for sex assigned at birth, estimated total intracranial volume (eTIV, from FreeSurfer) or total brain volume (from FSL), and socioeconomic status (SES) as defined by adjusted household income (total household income / number of household individuals) for the HCP cohort, or education level for the UK Biobank sample. Robust standard error estimates were calculated unless otherwise specified, and standardized β-coefficients are reported for all models along with Cohen’s *f* effect sizes for partial effects in multivariate regression.

In the HCP cohort, we investigated relationships between age, total cerebellar volume as defined by ACAPULCO parcellation (Han et al., 2020), total neocortical volume as defined by FreeSurfer *aparc* parcellation, eTIV as defined by FreeSurfer *aparc* parcellation, and MoCA total scores. Voxel-based morphometric (VBM) analyses considering years of age or MoCA scores as independent variables, corrected for eTIV, were conducted using the CAT12 toolbox (Gaser and Dahnke, 2016; https://neuro-jena.github.io/cat/), which utilizes a standardized pipeline adapted for VBM strategies in SPM12 (Ashburner and Friston, 2005) considering partial volume effects and bias correction of intensity non-uniformities. Statistical conjunction of corrected maps was conducted using FreeSurfer. Per Welch’s two-sample t-test, male and female HCP individuals differed in their neocortical volume [*t*(613·65)=−10·89, *p*<0·0001, male cortex mean volume: 478,998, female cortex mean volume: 438,824], cerebellar volume [*t*(651·48)=−11·99, *p*<0·0001, male cerebellar mean volume: 137,083, female cerebellar mean volume: 125,866], and eTIV [*t*(690·86)=−19·19, *p*<0·0001, male eTIV mean volume: 1,645,170, female eTIV mean volume: 1,425,052]. Females also had greater MoCA total scores [*t*(631·04)=2·79, *p*=0·005, male mean MoCA score: 26·01, female mean MoCA score: 26·55]. No sex differences were present for adjusted household income (*p*=0·097, male mean adjusted income: 68,943, female mean adjusted income: 59,750), or age (*p*=0·65, male mean age: 60·35, female mean age: 59·82).

In our UK Biobank validation sample, similar regression models were conducted to examine cerebellar volume in relation to age and cognitive performance on the DSST and TMB tasks. Gaussian mixture models (GMM, Bishop, 2006; McLachlan & Peel, 2000) were employed to classify participants into high and low cerebellar volume groups based on their average crus I and crus II volumes, allowing for more detailed subgroup analyses. In the ADNI cohort, we assessed relationships between cerebellar and neocortical volumes, age, APOE genotype, and MoCA scores. We also examined interaction effects between cerebellar volume and clinical diagnosis by amyloid status (high or low amyloid burden), age, and APOE allele zygosity, providing insights into the influence of genetic risk factors on cognitive performance.

## Results

### Age and cognition associations with neocortex and cerebellar volume

We found significant inverse relationships (**Figure 1a-b**) between age and both neocortical volume (β=−1,882.78, se=65.82, *p*<0·00001, Cohen’s *f*=1·31) and cerebellar volume (β=−298.78, se=25.28, *p*<0·00001, Cohen’s *f*=0·54). Age and eTIV did not show a significant association (β=−0·01, se=0·31, *p*=0·984, Cohen’s *f*=0·003), given the constancy in cranial size throughout the lifespan. Greater MoCA total scores showed significant associations with both greater neocortical volume (β=0·00001, se=0·000001, *p*=0·00001, Cohen’s *f*=0·22) and greater cerebellar volume (β=0·0001, se=0·00001, *p<*0·00001, Cohen’s *f*=0·24). In line with previous brain reserve research (van Loenhoud et al., 2018), we also observed a relationship between eTIV and cognitive performance (MoCA total scores: β=0·002, se=0·001, *p*=0·040, Cohen’s *f*=0·10) (**Figure 1d-e**).

**Figure 1.**
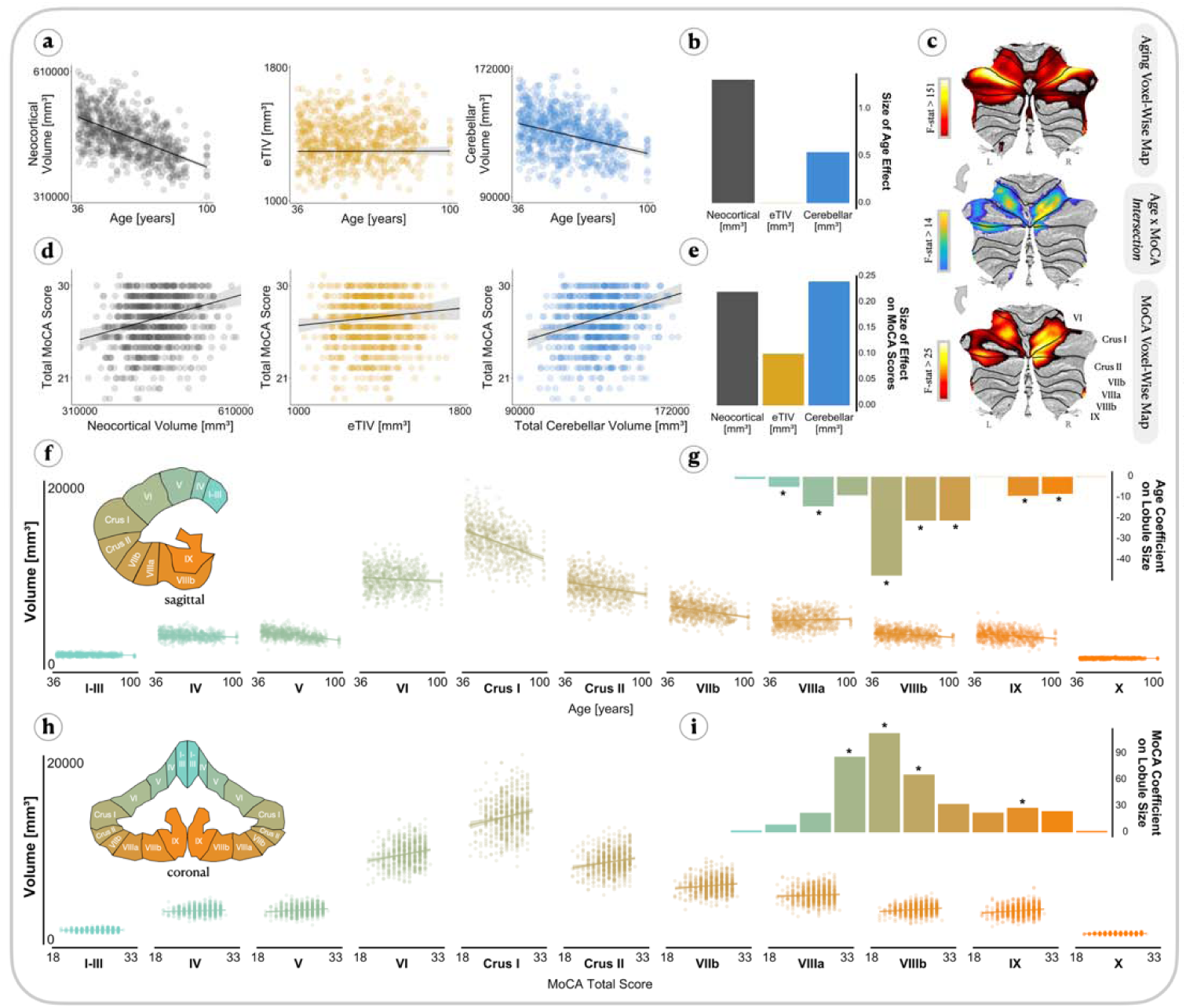
Aging-associated volume gradient across cerebellar lobules. (**a**) Correcting for sex assigned at birth, SES. and estimated total intracranial volume (eTIV), we found significant inverse relationships between age and neocortical volume (β=−1,882.78, se=−65.82. *p*<0.00001, Cohen’s *f*=1.31) and cerebellar volume (β=−298.78, se=23.28, *p*<0.00001, Cohen’s *f*=−0.54). Age and eTIV were not related (β=−0.01. se=0.31, *p*=0.984. Cohen’s *f*=0.003) The magnitude of these effects in relation to increasing age is displayed in (**b**). a plot of the Cohen’s *f* effect size for multivariate effects for age on neocortical volume. eTIV. and cerebellar volume, (**c**) <u>Top:</u> F-test association map of clusters in the cerebellum gray matter associated with years of age, correcting for eTIV (pFWE-corr<0.0001, qFDR=0.002, F<1,705), k=21). Brighter color (yellow) indicates greater years of age relating to lower cerebellar volume [mm^3^]. <u>Bottom:</u> F-test map of clusters in the cerebellum gray matter associated with MoCA Total scores, correcting for eTIV (pFWE-corr<0.0001, qFDR=0.002, F(1,705), k=21). Brighter color (yellow) indicates greater MoCA Total scores relating to greater cerebellar volume [mm^3^]. <u>Middle:</u> Conjunction map visualizing the intersection of the Aging vs MoCA voxel-wise result maps, suggesting that variations in this map contribute to aging-related changes in cognitive performance as measured by the MoCA. Crus I/Crus II region shows the strongest pattern of correlation between increasing age and a concomitant decrease in cerebellar volume and cognitive abilities, (**d**) Correcting for the same covariates as in (a), greater MoCA total scores related to greater neocortical volume (β=0.00001, se=0.000001, *p*=0.00001, Cohen’s *f*=0.22), and greater cerebellar volume (β=0.0001, se=0.00001, *p*<0.00001, Cohen’s *f*=0.24). We also observe a relationship between eTIV and cognition, MoCA total scores (β=0.002, se=0.001, *p*=0.040, Cohen’s *f*=0.10). The difference between neocortical, eTIV, and cerebellar volume in predicting cognitive scores is displayed in (**e**), showing the Cohen’s *f* effect sizes, on MoCA scores, (**f**) Aging-related effects on volume across cerebellar lobules demonstrates a striking gradient in the impact of aging on lobules. A subset cartoon of a sagittal plane cerebellum complements the scatterplot, offering a visual reference with a gradient mirroring the aging-related volume associations. (**g**) Correcting for sex, socioeconomic status (SES), and estimated total intracranial volume (eTIV), age was a significant predictor of volume for lobules IV, V, crus I, crus II, VIIb, VIIIb, and IX (bonferroni-corrected, alpha=0.05, *p*<0.05). This gradient underscores generally increasing aging-related changes from lobules I-III to Crus I and a decreasing gradient from Crus I to Lobule X. (**h**) MoCA-related effects on volume across cerebellar lobules demonstrates a striking gradient in the association of MoCA scores and cerebellar volume. A subset cartoon in the coronal plane of a cerebellum complements the scatterplot, offering a visual reference for a gradient mirroring the aging-related cerebellar volume associations, (**i**) Correcting for sex, socioeconomic status (SES), and eTIV, greater MoCA total scores were associated with higher volume in lobules VI, crus I. crus II, and VIIIb (bonferroni-corrected, alpha=0.05, *p*<0.05).

We next examined the above relationships using the individual ACAPULCO cerebellar lobule parcellations. The cerebellum does not age homogeneously, with age acting as a significant predictor of volume for lobules IV, V, crus I, crus II, lobule VIIb, VIIIb, and IX (Bonferroni-corrected, alpha=0·05, *p<*0·05) (**Figure 1f-g**). This differential aging forms a gradient, increasing from anterior lobules I-III to posterior lobe crus I, and decreasing from crus I to lobule X. Greater cerebellar volume values related to higher MoCA scores for lobules VI and VIIIb, and crus I and II (Bonferroni-corrected, alpha=0·05, *p<*0·05) (**Figure 1h-i**), exhibiting a similar mirrored gradient as in the age models.

In VBM analyses, years of age were associated with cerebellar voxel clusters along bilateral lobules V, VI, crus I, crus II (pFWE-corr<0·0001, qFDR=0·002, F(1,705), k=21) (**Figure 1c: top**). MoCA values showed associations along cerebellar voxels encompassing bilateral lobule V, VI, crus I, and crus II (pFWE-corr<0·0001, qFDR=0·002, F(1,705), k=21) (**Figure 1c: bottom**). To identify cerebellar cortex which shows both aging and cognitive effects, we conducted a statistical conjunction, where cerebellar voxels were kept in the cerebellar gray matter if they were present in both the aging and the MoCA maps (**Figure 1c: middle**).

In post-hoc supplementary analyses, we examined the voxel-wise relationship between cerebellar gray matter and subscales of the MoCA. These maps demonstrated both unique and shared representations between subscales and cerebellar gray matter (p-uncorr<0·001, **Figure S1**), with the MoCA Visuospatial subscale map surviving correction for multiple comparisons (pFWE-corr<0·0001, qFDR=0·002). The remaining maps did not survive correction for multiple comparisons, and were not further investigated.

**Figure S1.**
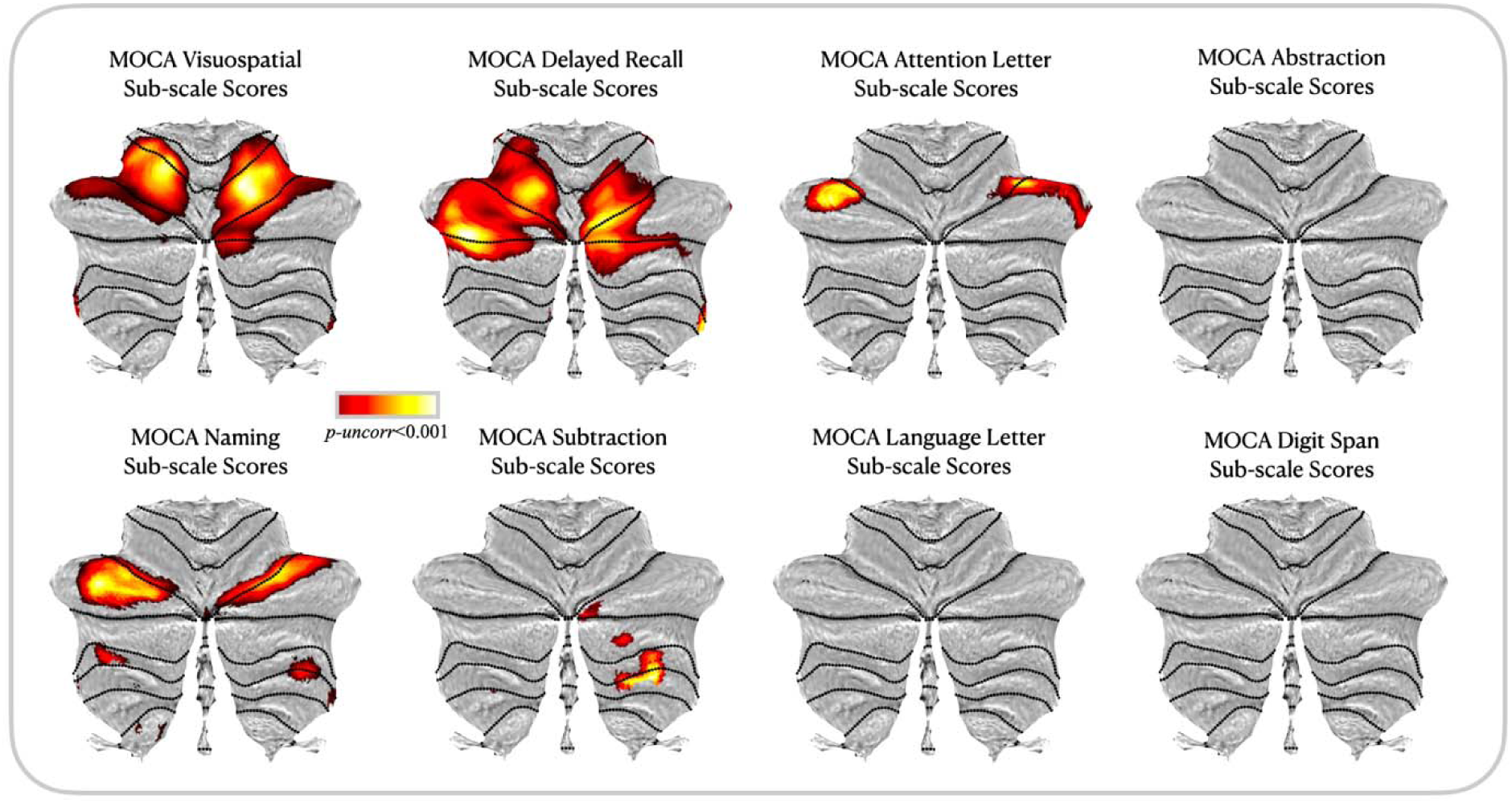
Cerebellar cognitive signatures: MoCA Sub-scale Score Maps. Voxel-wise cerebellar gray matter associations with MoCA scores from the visuospatial, delayed recall, language letter, abstraction, naming, mental subtraction, attention letter, and digit span subscales. Voxel-morphometric mapping demonstrates unique and shared representations between subscales and cerebellar gray matter, with the visuospatial subscale map surviving correction for multiple comparisons (pFWE-corr<0.0001, qFDR=0.002). The other subscale maps did not display a strong association between scores and cerebellar gray matter, and are visualized here with a *p*<0.001 threshold.

### Cognitive signatures in the cerebellum and their contribution to cognitive resiliency in healthy older adults

The spatial extent of the observed MoCA x Aging cerebellar signature (Aging-MoCA ROI) overlapped with underlying representations of the default mode and frontoparietal network as per the Buckner 7-Network parcellation (**Figure 2a: top left**), which maps neocortical resting-state functional networks to the cerebellum (Buckner et al., 2011), and attention, language, and motor functional representations as per the Multimodal Domain Task Battery (MDTB) parcellation (**Figure 2a: top right**), which maps functional task fMRI activity to cerebellar voxels (King et al., 2019). We conducted the same procedure for the Visuospatial MoCA x Aging conjunction map (Aging-Visuospatial ROI), and found that cerebellar clusters overlapped with voxels encompassing default mode, frontoparietal, ventral attention, and somatomotor networks as per the Buckner 7-Network parcellation (**Figure 2a: bottom left**), and attention, language, and motor functional representations as per the MDTB parcellation (**Figure 2a: bottom right**).

**Figure 2.**
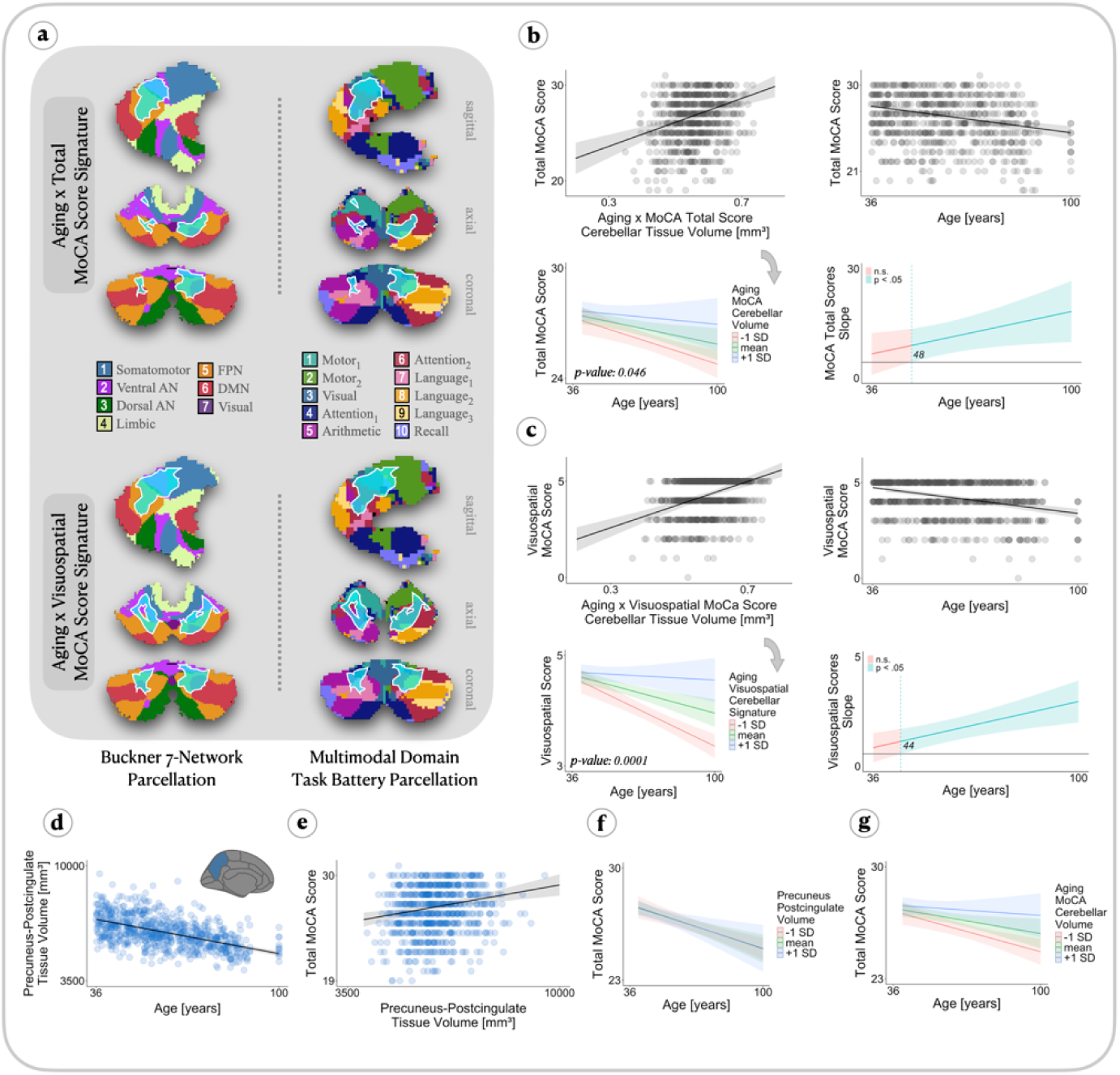
Cerebellar cognitive signature comprises regions related to cerebello-cerebral networks and acts as reserve factor against aging effect on cognitive performance. (**a**) Resulting conjunction intersection clusters of the Aging vs MoCA Total scores voxel-wise map (top) and Aging vs MoCA Visuospatial sub-scale scores map (bottom). The Aging-MoCA cluster (top) encompassed default mode and frontoparietal network representations as per the 7-Buckner cerebellar network parcellation (left), and attention, language, and motor functional representations as per the Multimodal Domain Task Battery parcellation (right). The Aging-Visuospatial cluster (bottom) encompassed default mode, frontoparietal, ventral attention, and somatomotor network representations (left), as well as attention, language, and motor functional networks (right). (**b-c**) Correcting for sex assigned at birth, SES, and eTIV, greater Aging-MoCA cerebellar volume (mm^3^) related to greater Total MoCA scores (β=12.50, se=2.12, *p*<0.0001, Cohen’s *f*=0.26) (**b:top left**), and greater Aging-Visuospatial scale volume (mm^3^) related to greater MoCA-Visuospatial scores (β=6.01, se=0.76, *p*<0.0001, Cohen’s f=0.37) (**c: top left**). Further, greater age related to lower MoCA total scores (β=−0.04, se=0.01, p<0.0001, Cohen’s *f*=0.28) (**b:top right**), and lower Aging-Visuospatial volume (β=−0-02, se=0.002, *p*<0.0001, Cohen’s *f*=0.37) (**c: top right**). The deleterious effect of age on MoCA scores was attenuated for individuals with greater levels of Aging-MoCA cerebellar volume (β=0.20, se=0.10, *p*<0.05, Cohen’s *f*=0.10) (**b: bottom left**), demonstrating a putative protective mechanism defining the MoCA cerebellar signature. The Aging-Visuospatial cerebellar signature also attenuated the effect of age on visuospatial scores (β=0.14, se=0.04, *p*=0.0001, Cohen’s *f*=0.15) (**c: bottom left**). Simple slopes analysis (pFDR<0.05), demonstrated that the protective effect of the Aging-MoCA cerebellar signature became most pronounced approximately after 48 years of age (**b: bottom right**), while the protective effect of the Aging-Visuospatial cerebellar signature on visuospatial scores became most pronounced approximately after 44 years of age (**c: bottom right**). (**d-g**) Post-hoc linear regression models, controlling for sex, eTIV, and SES, examined the relationship between precuneus-postcingulate cortex (PPCC) volume, aging, cognition, and cognitive reserve. (**d**) Age was inversely related to PPCC volumes (β=−30.25, se=1.52, Cohen’s *f*=0.88, *p*<0.0001), consistent with known neocortical aging patterns. While higher PPCC volume values were associated with greater total MoCA scores (β=0.001, se=0.0001, Cohen’s *f*=0.18. *p*=0.0001) (**e**), PPCC was not a significant moderator of the Age vs MoCA scores relationship (*p*=0.931) (**f**). (**g**) Inclusion of PPCC volume in the cerebellar reserve model did not alter the cerebellum’s unique role in mitigating age-related cognitive decline (*p*=0.033).

We extracted mean volume values from both the Aging-MoCA map, and the Aging-Visuospatial map. Aging-MoCA volume related to greater MoCA total score values (β=12·50, se=2·12, *p<*0·0001, Cohen’s *f*=0·26) (**Figure 2b: top left**). As expected from lobule-wide results in **Figure 1h**, we confirmed that age was related to lower MoCA total scores (β=−0·04, se=0·01, *p<*0·0001, Cohen’s *f*=0·28) (**Figure 2b: top right**). However, we can pinpoint the role of the cerebellum as a source of reserve by examining the moderation and visualizing participants according to their overall cerebellum size. The effect of age on MoCA total scores was attenuated in individuals with greater cerebellar volume in the Aging-MoCA intersection (β=0·20, se=0·10, *p*=0·046, Cohen’s *f*=0·10), suggesting a putative protective mechanism within this region of the cerebellum (**Figure 2b: bottom left**). We found similar patterns when investigating the effect of the Aging-Visuospatial ROI values on visuospatial subscale scores (β=6·01, se=0·76, *p*<0·0001, Cohen’s *f*=0·35) (**Figure 2c: top left**), as well as a negative effect of age on visuospatial subscale scores (β=−0·02, se=0·002, *p*<0·0001, Cohen’s *f*=0·37), (**Figure 2c: top right**). Further, Aging-Visuospatial ROI volume values also moderated the deleterious effect of age on visuospatial subscale scores (**Figure 2c: bottom left**), whereby individuals with greater cerebellar volume in the Aging-Visuospatial ROI demonstrated resilience against cognitive aging (β=0·14, se=0·04, *p*=0·0001, Cohen’s *f*=0·15). The protective effect for the Aging-MoCA cerebellar ROI became most pronounced and significant approximately after 48 years of age as per simple slopes analysis (pFDR<0·05), visualized with Johnson Neyman plots (**Figure 2b: bottom right**). Similarly, the Aging-Visuospatial ROI volume began to significantly confer protection against cognitive aging after 44 years of age (**Figure 2c: bottom right**).

Is the reserve effect specific to the cerebellum, or do the regions of neocortex that show connectivity with our cerebellar regions of interest also play a similar role? Based on previously reported structural and functional cerebello-cerebral networks, lower cerebellar lobules VI, crus I, and crus II show connectivity to posterior midline cerebral cortex (Wang et al., 2024; Xue et al., 2021; Buckner et al., 2011). We thus created a volume meta-ROI from FreeSurfer bilateral precuneus and post-cingulate cortex (PPCC). We used this ROI in supplementary post-hoc linear regression models controlling for the same covariates as earlier. Higher volume values in the PPCC ROI were associated with higher volume values in the Aging-MoCA ROI (β=6,376, se=559, *p*<0·0001, Cohen’s *f*=0·50). While age was inversely related to PPCC volume (β=−30·25, se=1·52, *p*<0·0001, Cohen’s *f*=0·88) (**Figure 2d**), and higher PPCC volume was associated with greater MoCA scores (β=0·001, se=0·0001, *p*=0·0001, Cohen’s *f*=0·18) (**Figure 2e**), we surprisingly find that PPCC volume did not act as a moderator of the inverse age vs MoCA relationship (*p*=0.931) (**Figure 2f**). We further confirmed the specificity of the brain reserve effect to the cerebellum by including the PPCC ROI in our Aging-MoCA cerebellar model (MoCA scores ~ MoCA cerebellar volume x Age + PPCC volume). The Aging-MoCA cerebellar ROI continued to significantly mitigate the inverse age vs MoCA score association (*p*=0.033) (**Figure 2g**).

### Generalizability and specificity of cerebellar reserve to independent cognitive tasks

To examine whether the protective effect from the Aging MoCA-associated cerebellum would generalize to independent tasks in HCP participants, we examined four tasks for which we had available data: the Dimensional Card Sorting Task (Gur et al., 2001), and the Flanker Attention Task (Zelazo, et al., 2013), which engage the frontoparietal network (Lie et al., 2006; Fassbender, et al., 2006), and two tasks that require memory processes, namely the Picture Sequence Memory Task (Weintraub et al., 2013), and the List Sorting Working Memory Task (Gershon, et al., 2013). In linear interaction models controlling for sex and eTIV, we found that greater volume in the Aging-MoCA cerebellar ROI conferred a protective effect on the Card Sorting task (β=0·96, se=0·35, *p*=0·007, Cohen’s *f* effect size=0·11) (**Figure 3a: left**), and the Flanker Attention Task (β=0·77, se=0·32, *p*=0·016, Cohen’s *f* effect size=0·10) (**Figure 3b: left**), such that age-related declines in these cognitive task scores were mitigated in individuals with greater Aging-MoCA ROI volume. As per simple slopes analysis (pFDR<0·05), the association between MoCA-related cerebellar volume and cognitive scores was more pronounced in individuals older than 55 years for both the Card Sorting Task (**Figure 3a: right**) and the Flanker Attention Task (**Figure 3b: right**). We did not find a significant protective effect of the Aging-MoCA ROI for either the Picture Memory Sequence Task (*p*=0·64) (**Figure 3c**) or the List Sorting Working Memory Task (*p*=0·71) (**Figure 3d**), consistent with this cerebellar ROI’s overlap with frontoparietal and not memory-related representations.

**Figure 3.**
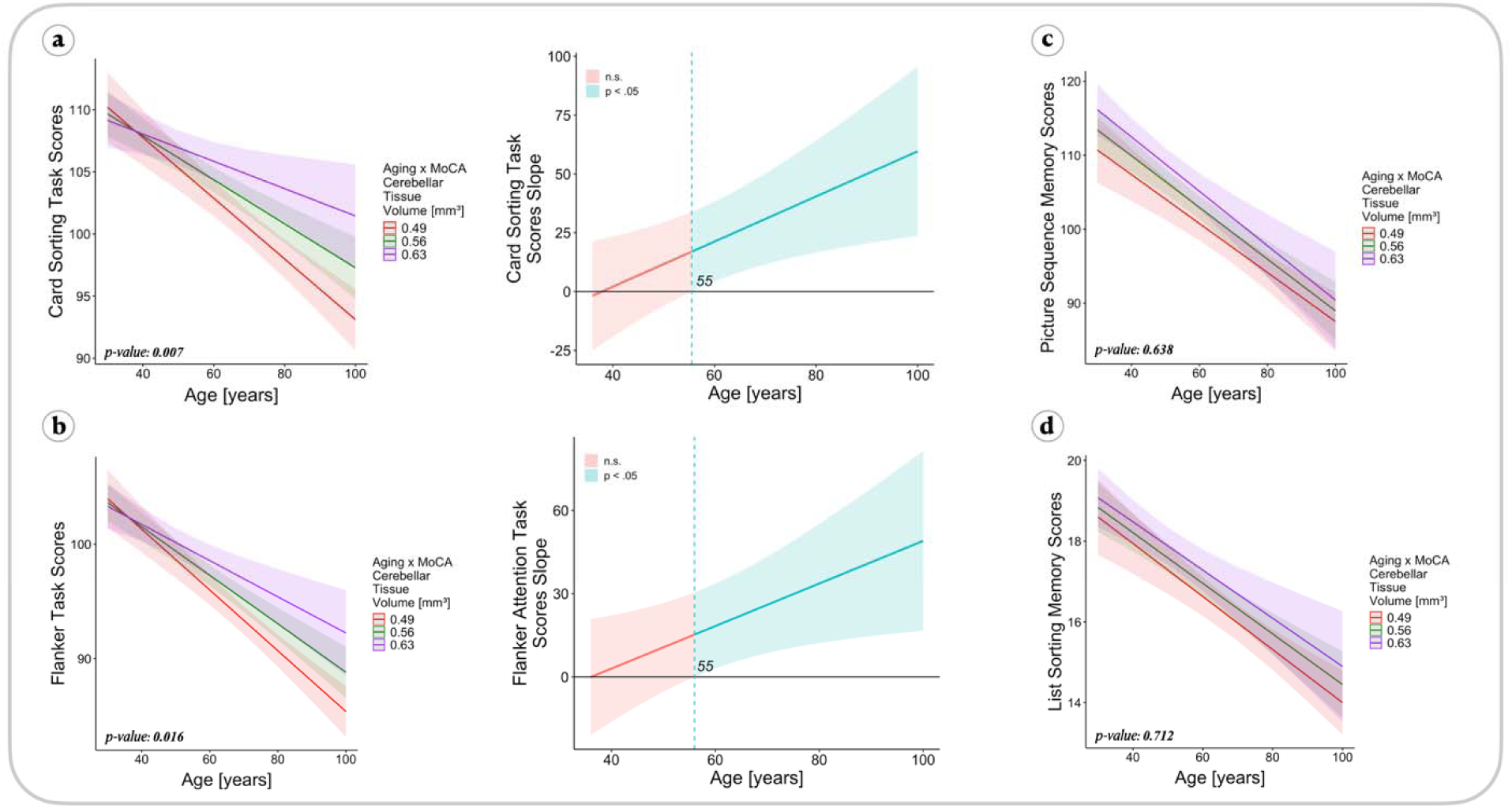
Greater rates of cerebellar MoCA signature help attenuate aging in a separate subset of tasks known to engage frontoparietal circuits. Controlling for sex and estimated intracranial volume in linear regression models, we found evidence that the MoCA-related cerebellar volume signature conferred a protective effect on other behavioral tasks known to engage frontoparietal network circuits, including the Dimensional Change Card Sorting task (β=0.96, se=0.35, *p*=0.007, Cohen’s *f*=0.11) (**a: left**), and the Flanker Attention Task (β=0.77, se=0.32, *p*=0.016, Cohen’s *f*=0.10) (**b: left**), tasks implicated in cognitive flexibility and executive function, and cognitive control and attention, respectively. Johnson Neyman analysis (pFDR<0.05) revealed that the association between cerebellar MoCA signature volume and cognitive scores was more pronounced in older individuals: for Card Sorting scores (significant cerebellar reserve effect after 55 years old) (**a: right**), and Flanker Attention Task scores (significant cerebellar reserve effect after age 55 years old) (**b: right**). These findings demonstrate that the MoCA-related cerebellar volumetric signature can promote a safeguard of cognitive resilience in related but separate cognitive tasks. We did not find interactions between cerebellar MoCA signature volume and age for the Picture Memory Sequence Task (**c**) or List Sorting Working Memory Task (**d**) (*p*>0.6). This overall suggests that individuals with greater cerebellar reserve may be better able to maintain these cognitive abilities in older age than those with lower cerebellar reserve, and that tasks emphasizing memory testing may be more involved with reserve in other regions of the cerebellum or cortex.

### Cerebellar reserve effect in the UK Biobank

We next sought to replicate our findings with an independent, large dataset of healthy older adults from the UK Biobank. With available volumetric data from crus I, crus II, and lobule IX, we first examined the relationship between age and cerebellar volume. Given HCP-Aging data (**Figure 1f**), we had the *a priori* hypothesis that the posterior lobule crus I would show the greatest aging effect, followed by crus II and lobule IX. We indeed found that the effect of age on cerebellar volume was most pronounced for crus I (β=−0·002, se<0·001, *p*<0·0001, Cohen’s *f* effect size=0·38), followed by crus II (β=−0·002, se<0·001, *p*<0·0001, Cohen’s *f* effect size=0·27), and lastly lobule IX (β=−0·003, se<0·001, *p*<0·0001, Cohen’s *f* effect size=0·16) (**Figure 4a-b**), mirroring and replicating the age-related gradient we identified in our previous analysis (**Figure 1f-g**).

**Figure 4.**
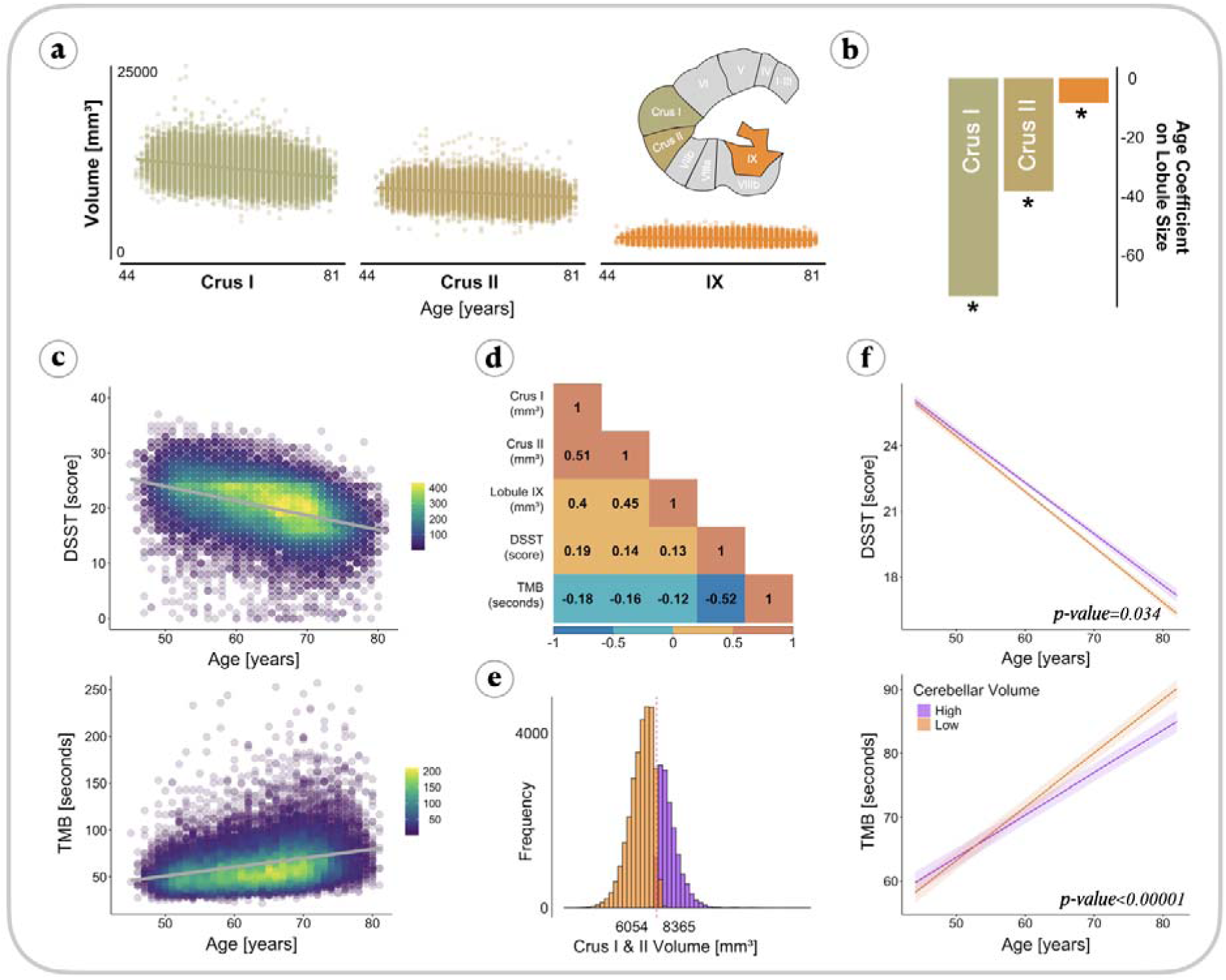
Cerebellum reproducibly modulates aging-related cognitive decline in the UK Biobank. Cerebellar regions exhibit varying aging-related effects, with Crus I showing the most pronounced age association. Utilizing a Gaussian mixture model (GMM) approach, we identify distinct cerebellar volume groups and demonstrate that individuals with high cerebellar volume experience attenuated aging-related declines in cognitive performance. **(a)** Replication of the HCP dataset reveals a decreasing gradient of age effects across three lobules—Crus I, Crus II, and Lobule IX, in robust linear models corrected for sex assigned at birth, total head volume, and socioeconomic status (SES). **(b)** Visual representation of the age effect magnitudes from Crus I, Crus II, and Lobule IX. Notably, Crus I exhibits a substantial aging-related decline, with a 60 unit volume decrease per year, double that of Crus II and Lobule IX. **(c)** Significant lower Digit Symbol Substitution Test (DSST) scores and greater time to complete Trails-Making 8 (TMB) with increasing age, corrected for same covariates. **(d)** Unadjusted Pearson correlations highlight strong associations between lobules (Crus I vs Crus II and Lobule IX, *r*>0.4) and between cognitive batteries (*r*>0.1). **(e)** Application of a GMM identifies two distinct distributions in a cerebellar Crus I and Crus II composite, enabling the classification of low and high cerebellar volume groups. **(f)** Exploration of cerebellar volume as a moderator in interaction models, correcting for sex, SES, and total head volume, reveals that individuals with high cerebellar volume experience less DSST (β=−0.02, se=0.01, *p*=0.034) and TMB (β=0.18, se=0.04, *p*<0.00001) age-related decline compared to those with lower cerebellar volume.

We then examined the relationship between age and cognitive scores from two tasks available to us, namely the DSST, and the TMB. Increasing age conferred lower DSST scores (β=−0·25, se=0·004, *p*<0·0001, Cohen’s *f* effect size=0·41) and longer TMB completion times (β=−0·81, se=0·02, *p*<0·0001, Cohen’s *f* effect size=0·31) (**Figure 4c**). Unadjusted correlations showed that crus I displayed the strongest correlation with tasks (crus I vs DSST: *r*=0·19, crus I vs TMB: *r*=−0·18), followed by crus II (crus I vs DSST: *r*=0·14, crus I vs TMB: *r*=−0·16), and finally lobule IX (crus I vs DSST: *r*=0·19, crus I vs TMB: *r*=0·14) (**Figure 4d**).

We next sought to replicate the effect that differences in cerebellar volume are associated with different cognitive trajectories. We found that individuals with high cerebellar volumes experienced mitigated rates of cognitive decline in both the DSST (β=−0·02, se=0·01, *p*=0·034) and TMB (β=0·18, se=0·04, *p*<0·00001) (**Figure 4e**). That is, at the age of 80, high cerebellar volume individuals were about 9 seconds faster to complete the Trails Making-B task and perform 5% better on the DSST. These findings replicated our cerebellar reserve models in the HCP data, now in a mega-dataset of older adults in the UK.

### Cerebellar reserve in the Alzheimer’s disease neuroimaging cohort

We next asked, if the cerebellum plays a role in healthy aging, might it also have a protective effect in clinical cognitive impairment and dementia? Examining the distribution of MoCA total scores by diagnosis showed that MoCA scores tended to decrease with each clinical diagnostic stage (F(2)=289·60, *p*<0·0001) (**Figure S2a**), with Tukey HSD post-hoc tests showing that all three groups differed significantly from one another (mean MoCA score in CN: 25.07; mean MoCA score in MCI: 22.36; mean MoCA score in AD: 16.99). Total neocortical volume showed concomitant decreases with clinical stage (F(2)=27·68, *p*<0·0001) (**Figure S2b**), with post-hoc tests showing significant differences between AD and MCI, and AD and CN, but not between CN and MCI groups (mean cortex volume in CN: 427,996.10; mean cortex volume in MCI: 423,554.40; mean cortex volume in AD: 395,070.80). Interestingly, cerebellar volume remained consistent across diagnostic categories (F(2)=0·74, *p*=0·475) (**Figure S2c**) (mean cerebellar volume in CN: 127,668; mean cerebellar volume in MCI: 128,135; mean cerebellar volume in AD: 126,778).

**Figure S2.**
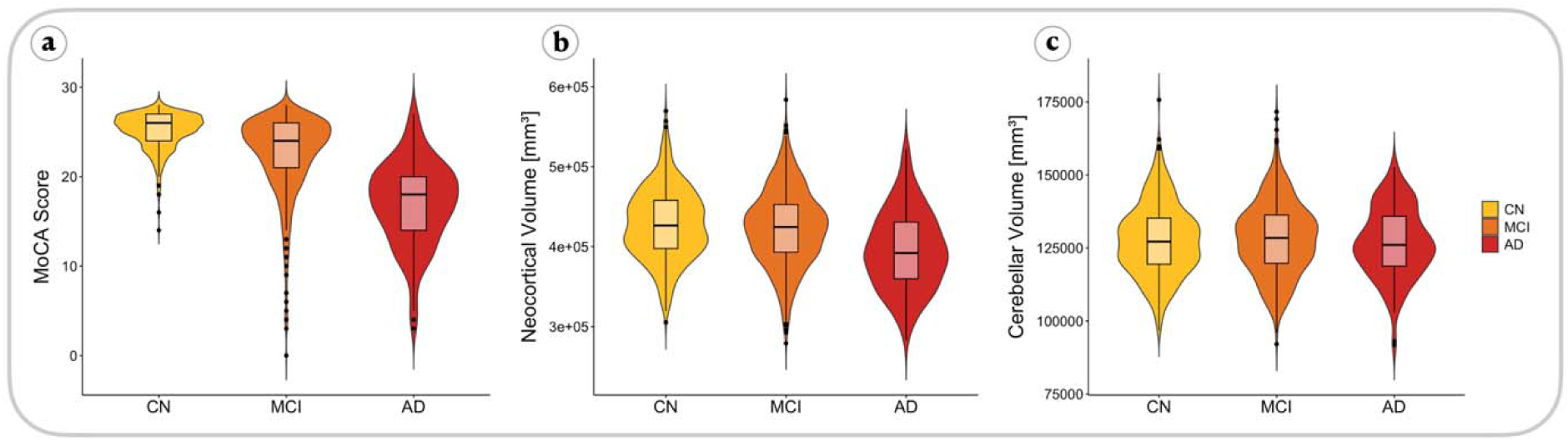
Decreasing MoCA scores and cortex volume by clinical diagnosis. Distributions of (**a**) MoCA scores by clinical diagnosis amongst clinically normal (CN), mild cognitive impairment (MCl), and Alzheimer’s disease (AD) showed significant differences as per analysis of variance (ANOVA) (F(2)=289.60, *p*<0.0001), with Turkey HSD post-hoc tests demonstrating that all three groups differed significantly from one another (mean MoCA score in eN: 25.07; mean MoCA score in MCl: 22.36; mean MoCA score in AD: 16.99). (**b**) Neocortical volumes also showed a significant effect of diagnosis group (F(2)=27.68, *p*<0.0001), with post-hoc tests showing significant differences between AD and MCI, and AD and CN, but not between CN and MCl groups (mean cortex volume in CN: 427,996.10; mean cortex volume in Mel: 423,554.40; mean cortex volume in AD: 395,070.80). (**c**) Comparing cerebellar volumes showed no significant effect of diagnosis (F(2)=0.74, *p*=0.475), (mean cerebellar volume in CN: 127,668.20; mean cerebellar volume in MCl: 128,135.30; mean cerebellar volume in AD: 126,777.90).

We next examined cerebellar volume relationships between MoCA scores and diagnostic group by amyloid burden (MoCA ~ cerebellar volume x diagnosis). We found a significant interaction between diagnosis and cerebellar volume in predicting MoCA scores for the amyloid-beta negative (Aβ−) AD group (β=0·0001, se=0·00001, *p*=0·0001) (**Figure 5a**) (**Table 2**). Three-way interaction models of age x cerebellar volume x diagnosis, predicting MoCA scores, correcting for the same covariates as in previous models, showed a trending age moderation effect on MoCA scores for the AD Aβ− group, and a moderation in AD Aβ+ (age x cerebellar volume x AD Aβ−: β=0·00001, se=0·000001, *p*=0·128; age x cerebellar volume x AD Aβ+: β=−0·00001, se=0·000001, *p*=0·036) (**Figure 5b**) (**Table 3**). Taken together, increasing age and lower cerebellar volume was associated with lower MoCA scores before substantial amyloid burden, whereas once amyloid burden becomes substantial, greater cerebellar volumes confer better MoCA scores only in adults who have yet to putatively accumulate other age-related brain changes and pathology.

**Figure 5.**
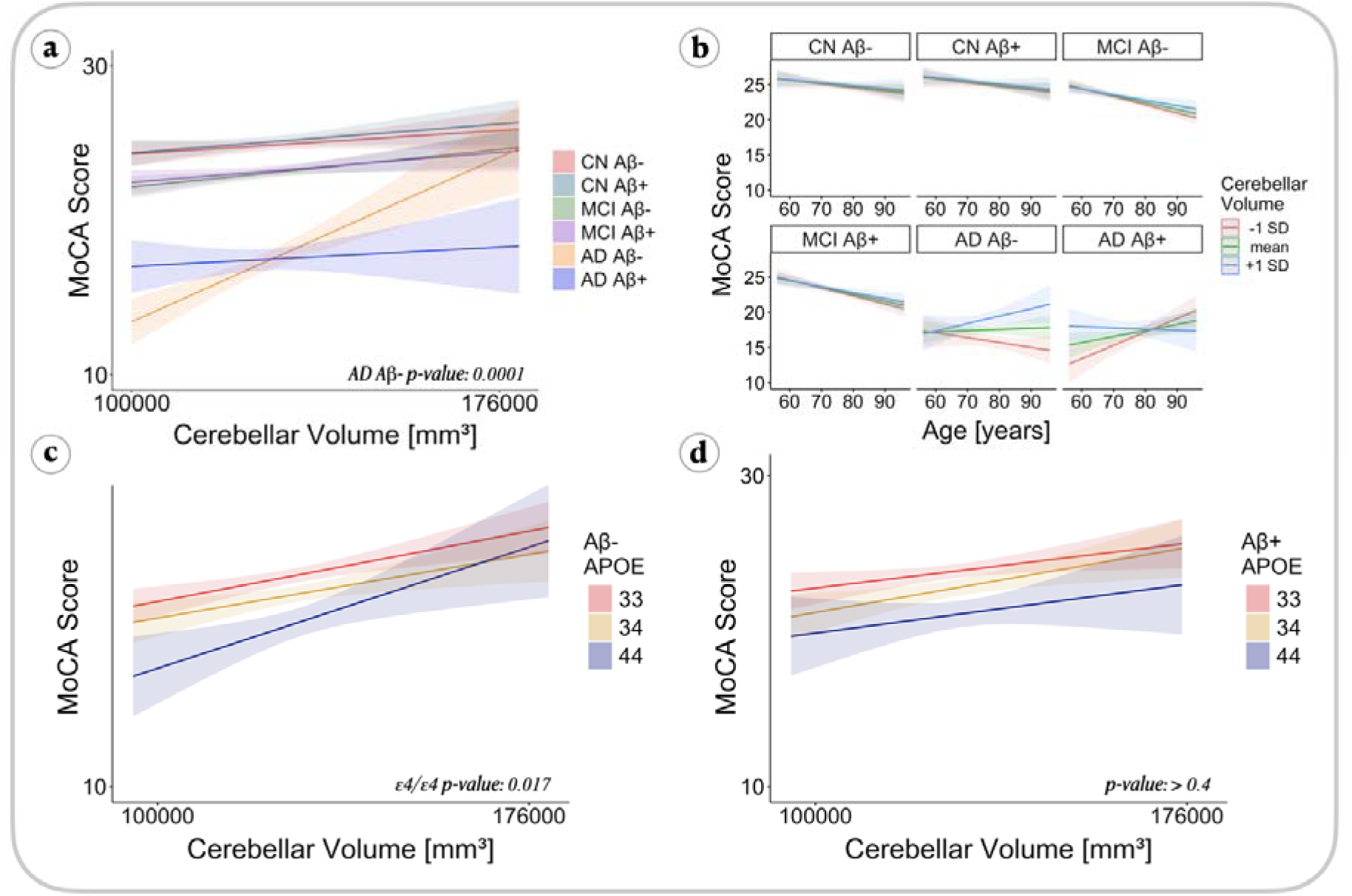
Cerebellar volume and MoCA Score relationships in ADNI. MoCA and cerebellar volume couple most in AD and APOEe4 prior to amyloidosis. In the Alzheimer’s Disease Neuroimaging Initiative (ADNI) cross-sectional cohort, we assessed the relationship between cerebellar total volume and total MoCA scores as a function of diagnosis and APOE gene zygosity in models correcting for sex and estimated intracranial volume (eTIV). (**a**) Amyloid-negative (Aβ−) individuals with an AD clinical diagnosis experienced the most pronounced relationship between cerebellar total volume and MoCA total scores (*p*=0.0001). (**b**) Investigating the effect of years of age on this relationship in 3-way interaction models (age x cerebellar volume x diagnosis), we found a trend in that age moderated the effects in Aβ− AD individuals such that greater cerebellar volume conferred attenuated cognitive decline in MoCA scores with increasing age (*p*=0.128), and a Significant age moderation effect in Aβ+ AD individuals who were younger than their counterparts (*p*=0.0.36), suggesting that cerebellar volume is most protective in individuals prior to the synergistic deleterious effects of increasing age and amyloid brain burden. (**c**) Stratifying individuals by APOE allele type across diagnostic groups, we found that in Aβ−individuals, cerebellar volume was most related to MOCA scores in the ε4/ε4 carriers (*p*=0.017), with no significant effects of APOE genotype found for Aβ+ individuals (**d**).

**Table 2:**
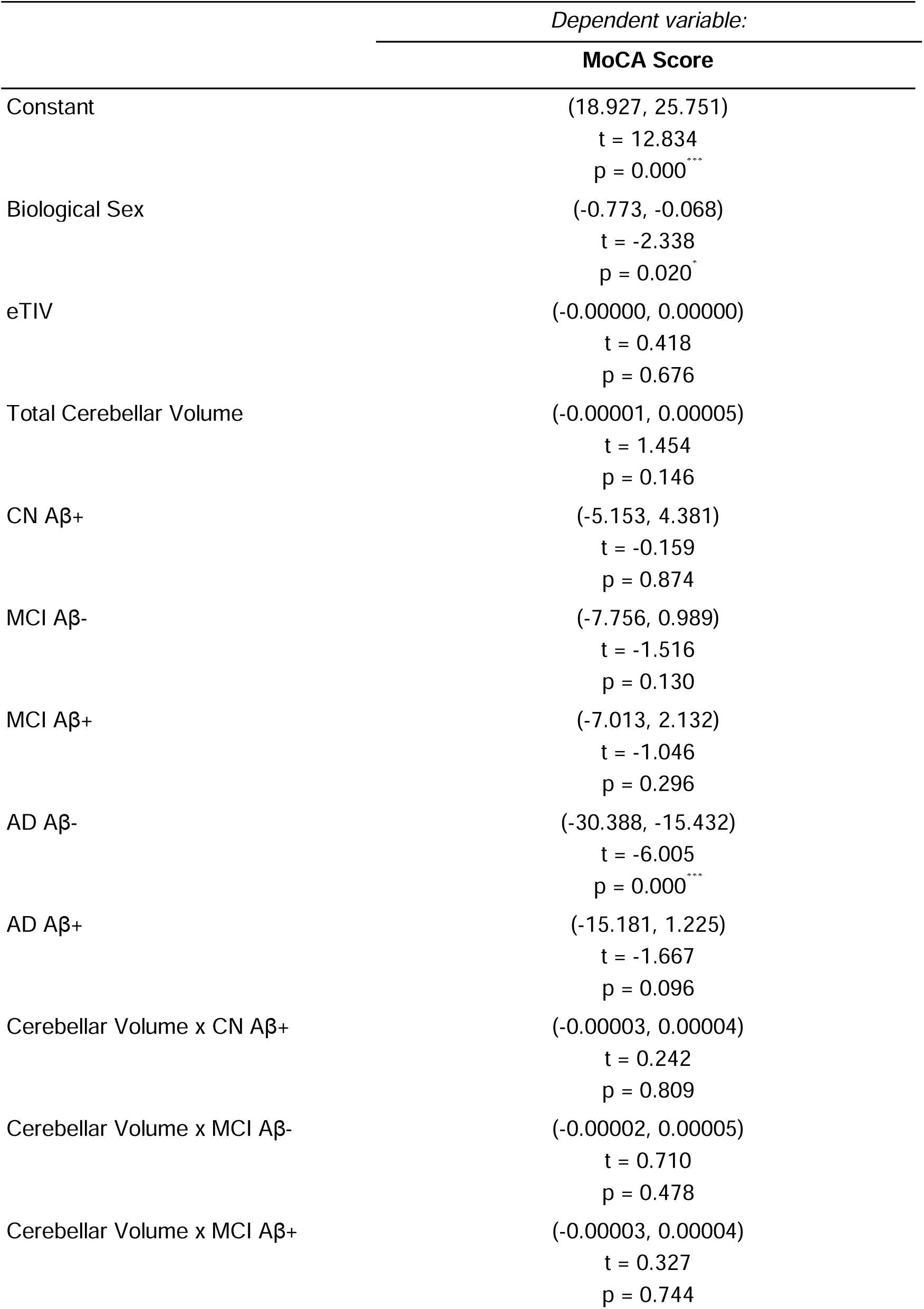
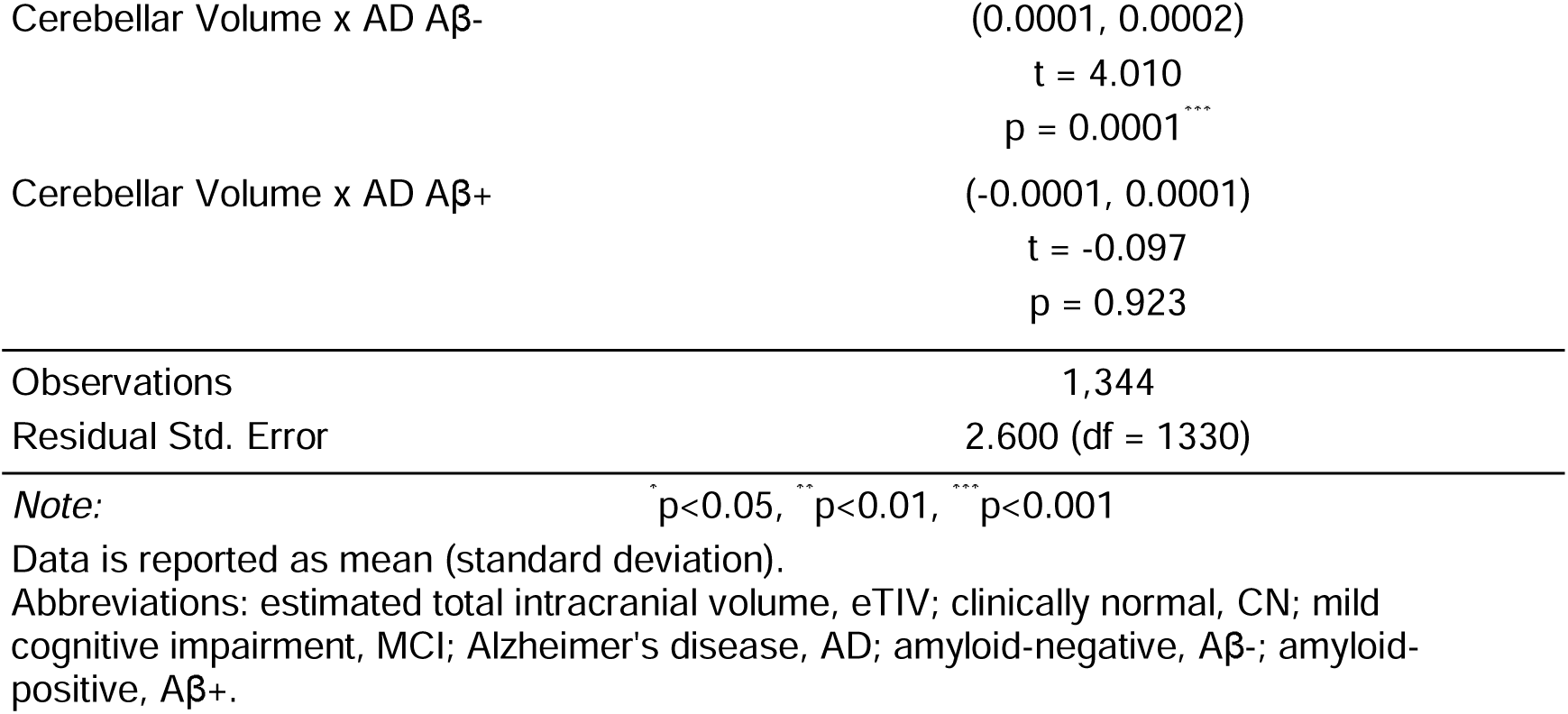
Cerebellar Volume x Diagnosis Aβ Group.

**Table 3:**
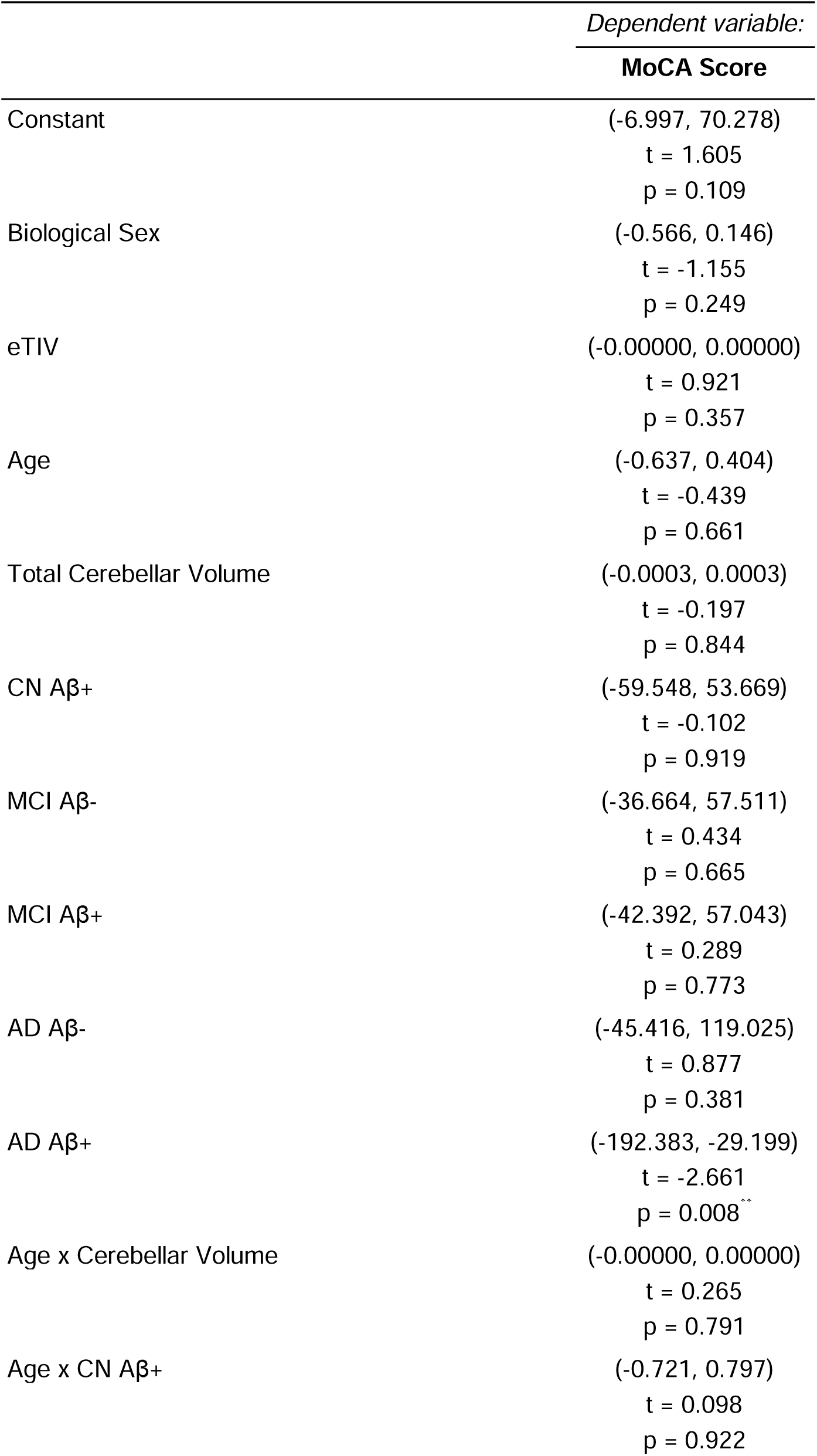
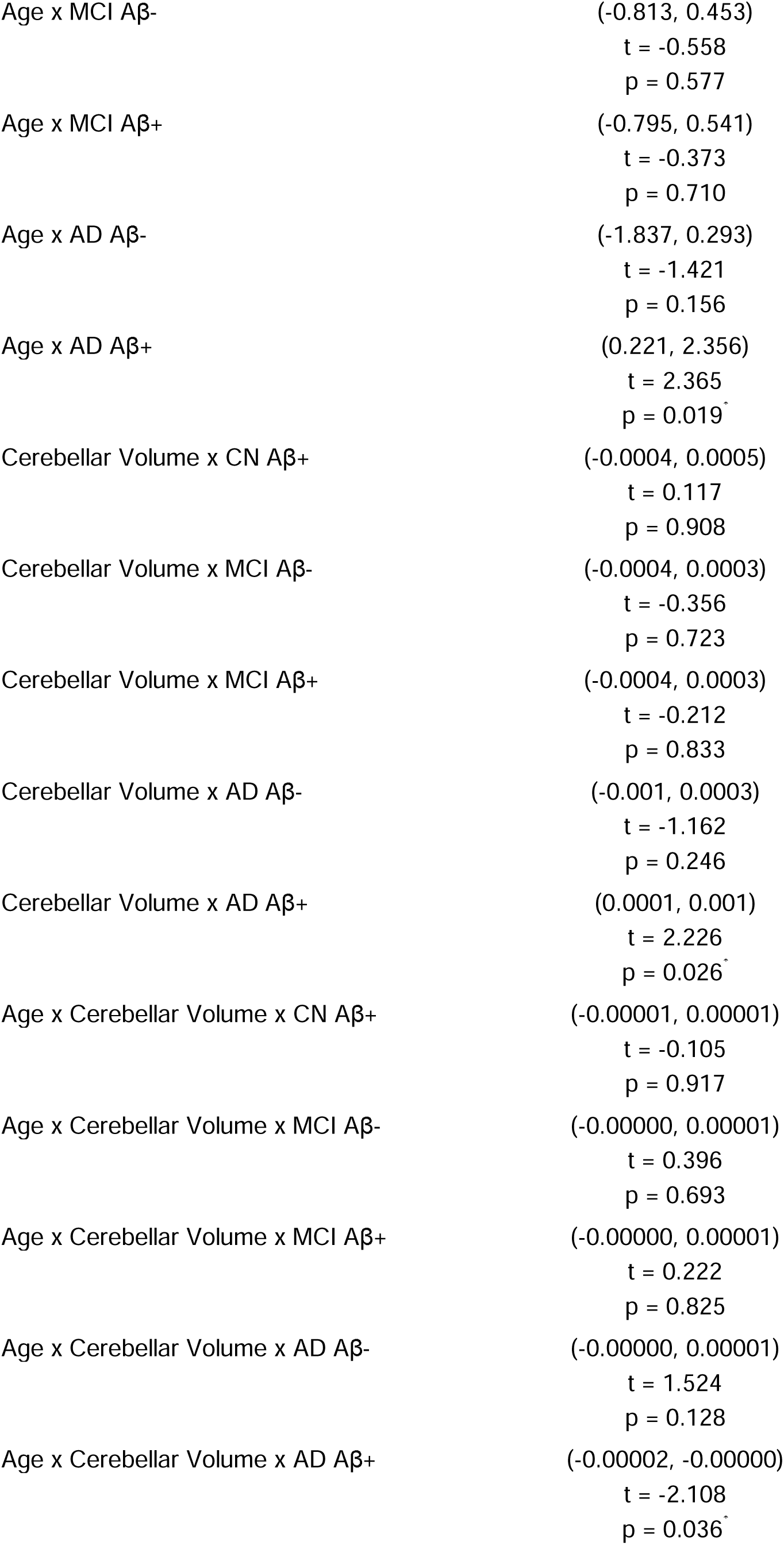
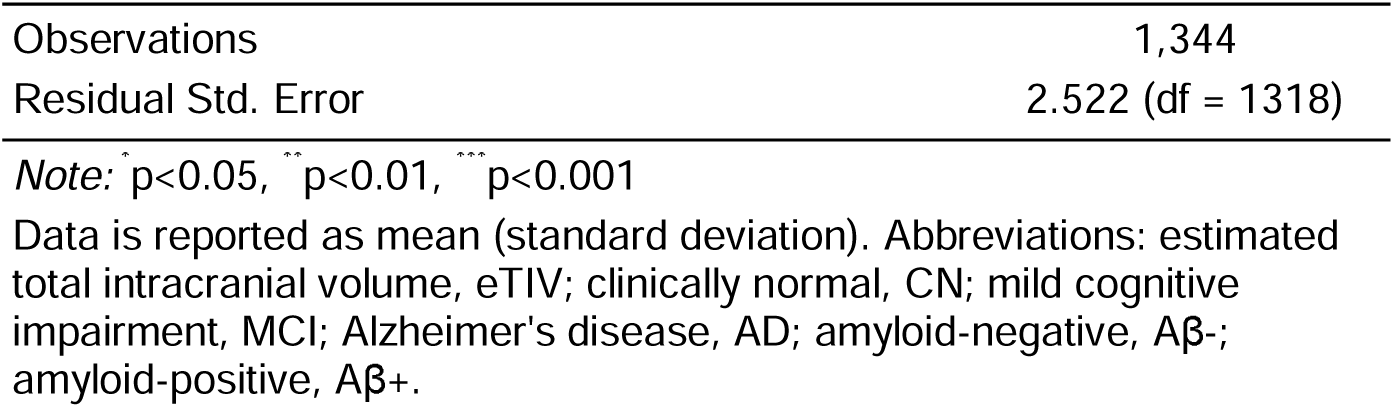
Cerebellar Volume x Age x Diagnosis Aβ Group.

|  | *Dependent variable:* |
| --- | --- |
|  | **MoCA Score** |
| Constant | (-6.997, 70.278)<br>t = 1.605<br>p = 0.109 |
| Biological Sex | (-0.566, 0.146)<br>t = -1.155<br>p = 0.249 |
| eTIV | (-0.00000, 0.00000)<br>t = 0.921<br>p = 0.357 |
| Age | (-0.637, 0.404)<br>t = -0.439<br>p = 0.661 |
| Total Cerebellar Volume | (-0.0003, 0.0003)<br>t = -0.197<br>p = 0.844 |
| CN Aβ+ | (-59.548, 53.669)<br>t = -0.102<br>p = 0.919 |
| MCI Aβ- | (-36.664, 57.511)<br>t = 0.434<br>p = 0.665 |
| MCI Aβ+ | (-42.392, 57.043)<br>t = 0.289<br>p = 0.773 |
| AD Aβ- | (-45.416, 119.025)<br>t = 0.877<br>p = 0.381 |
| AD Aβ+ | (-192.383, -29.199)<br>t = -2.661<br>p = 0.008** |
| Age x Cerebellar Volume | (-0.00000, 0.00000)<br>t = 0.265<br>p = 0.791 |
| Age x CN Aβ+ | (-0.721, 0.797)<br>t = 0.098<br>p = 0.922 |
| Age x MCI Aβ- | (-0.813, 0.453)<br>t = -0.558<br>p = 0.577 |
| Age x MCI Aβ+ | (-0.795, 0.541)<br>t = -0.373<br>p = 0.710 |
| Age x AD Aβ- | (-1.837, 0.293)<br>t = -1.421<br>p = 0.156 |
| Age x AD Aβ+ | (0.221, 2.356)<br>t = 2.365<br>p = 0.019<sup>*</sup> |
| Cerebellar Volume x CN Aβ+ | (-0.0004, 0.0005)<br>t = 0.117<br>p = 0.908 |
| Cerebellar Volume x MCI Aβ- | (-0.0004, 0.0003)<br>t = -0.356<br>p = 0.723 |
| Cerebellar Volume x MCI Aβ+ | (-0.0004, 0.0003)<br>t = -0.212<br>p = 0.833 |
| Cerebellar Volume x AD Aβ- | (-0.001, 0.0003)<br>t = -1.162<br>p = 0.246 |
| Cerebellar Volume x AD Aβ+ | (0.0001, 0.001)<br>t = 2.226<br>p = 0.026<sup>*</sup> |
| Age x Cerebellar Volume x CN Aβ+ | (-0.00001, 0.00001)<br>t = -0.105<br>p = 0.917 |
| Age x Cerebellar Volume x MCI Aβ- | (-0.00000, 0.00001)<br>t = 0.396<br>p = 0.693 |
| Age x Cerebellar Volume x MCI Aβ+ | (-0.00000, 0.00001)<br>t = 0.222<br>p = 0.825 |
| Age x Cerebellar Volume x AD Aβ- | (-0.00000, 0.00001)<br>t = 1.524<br>p = 0.128 |
| Age x Cerebellar Volume x AD Aβ+ | (-0.00002, -0.00000)<br>t = -2.108<br>p = 0.036<sup>*</sup> |
| Observations | 1,344 |
| Residual Std. Error | 2.522 (df = 1318) |

*Note:* ∼ $p < 0.05$ , $\sim\sim$ $p < 0.01$ , $\sim\sim\sim$ $p < 0.001$
Data is reported as mean (standard deviation). Abbreviations: estimated total intracranial volume, eTIV; clinically normal, CN; mild cognitive impairment, MCI; Alzheimer's disease, AD; amyloid-negative, Aβ -; amyloid-positive, Aβ +

Lastly, investigating moderation effects between cerebellar volume and APOE allele type on MoCA scores in Aβ− individuals (excluding ε2/ε2 carriers due to low numbers of individuals), we found that the relationship between cerebellar volume and MoCA scores was significant for ε4/ε4 homozygous carriers (**Figure 5c**) (cerebellar volume x APOE ε4/ε4: β=0·0001, *t*=2.39, se<0·0001, *p*=0·017), with trending relationships for ε3/ε3 carriers (β=0·0001, *t*=1·94, se<0·0001, *p*=0·052), and ε3/ε4 carriers (β=0·0001, *t*=1·75, se<0·0001, *p*=0·081). No significant effect of zygosity on cerebellar x MoCA scores was found for Aβ+ individuals (*p*>0·4) (**Figure 5d**). In post-hoc supplementary models, we did not find any significant interaction of age for any APOE allele combination (MoCA ~ age x APOE) in either Aβ− (*p*>0·2), or Aβ+ (*p*>0·1) individuals.

Together, these analyses demonstrated that cerebellar volume relationships with MoCA scores are most relevant for Aβ− individuals with the greatest genetic risk for AD, those being ε4/ε4 APOE carriers who have not yet progressed past a putative resilience threshold designated by Aβ burden status, extending the threshold model of brain reserve (Satz, 1993; Stern et al., 2019) to include the cerebellum.

## Discussion

In this cross-sectional study, we investigated cerebellar structural integrity and its relation to cognitive aging across different cohorts. Our spatial analysis of cerebellar structure revealed that the cerebellum ages differentially across the anterior-posterior axis. Aging effects are most pronounced in the posterior cerebellum, particularly crus I, relative to more anterior regions like lobules I-III (**Figure 1**), providing a more complete portrait of its structural aging which aligns with previous research documenting reductions in cerebellar structure (Raz et al., 2005; Bernard et al., 2015). We further found that higher cognitive scores from the MoCA testing battery were associated with greater tissue density in the cerebellum, especially in areas of cerebellar cortex that show connectivity to functional cerebral networks such as the frontoparietal, default mode, and ventral attention networks (**Figure 2**). Notably, greater volume along the posterior lobe conferred a protective effect against age-related decline in MoCA performance. We validated this finding in a large, independent dataset using tasks of processing speed and executive function from the UK Biobank (**Figure 4**). These results are consistent with previous studies generally implicating lobules VI, crus I, and crus II in cognitive performance such as processing speed and visual reproduction (Abe et al., 2008; Lee et al., 2005).

The protective influence of MoCA-related cerebellar tissue extended beyond general cognitive performance to tasks that engage intrinsic frontoparietal networks (**Figure 3**). The consistency of our findings across separate cohorts provides strong evidence for a cerebellar reserve in cognitive aging. That is, individuals with larger posterior cerebella show significantly decreased cognitive decline, while those with below-average volumes demonstrate relatively accelerated trajectories. In addition to its replicability, we find that the reserve effect seemed specific to the cerebellum. We tested whether tissue density in neocortex, which provides a large descending pathway via corticopontine pathways (Ramnani et al., 2006; Leiner, Leiner and Dow, 1993), would confer some level of protection against cognitive aging. We found that cerebellar volume carried a unique contribution to preserving cognition above and beyond cerebral cortex (**Figure 2**). This suggests that cerebellar volume may serve as a unique biomarker for cognitive resilience, potentially delaying the identification of at-risk individuals by masking overt clinical symptoms.

In the ADNI study, cerebellar volume showed the strongest associations with cognition in patients with a clinical diagnosis of AD and low amyloid-beta brain burden, particularly in APOE ε4/ε4 carriers (**Figure 5**). Taken together, our findings underscore a potentially large role for the cerebellum in cognitive resilience, and fall in line with the threshold model of brain reserve (Satz, 1993; and Stern et al., 2019 for a detailed discussion), now extending this model to include the cerebellum. This study overall highlights the cerebellum as a modulator of cognitive aging, and its potential as a therapeutic target for interventions for cognitive impairment, especially in populations at greater genetic risk for AD.

Our analysis in the ADNI cohort further revealed relatively stable cerebellar structure across different clinical stages (**Figure S2**), in contrast to the pronounced neocortical atrophy found in advanced stages along the AD trajectory (Dickerson et al., 2009). This suggests a certain robustness for the cerebellum to disease-related atrophy, relative to neocortical structures, and positions it to serve as a source of cognitive reserve in AD and cognitive decline (Arleo et al., 2024; Jacobs et al., 2018). Further, cerebellar volume showed the strongest associations with cognition in patients with a clinical diagnosis of AD dementia and low amyloid burden, particularly in APOE ε4/ε4 carriers (**Figure 5**). These findings overall suggest that cerebellar reserve mechanisms may be particularly important in the early stages of cognitive decline before substantial accumulation of amyloid pathology.

Despite its large sample size and replicability across study cohorts, several limitations must be acknowledged. The cross-sectional nature of our analysis limits our ability to draw causal conclusions about the temporal progression of the relationship between cerebellar atrophy and cognitive decline. Longitudinal analyses are needed to clarify the role of the cerebellum in the progression of cognitive impairment. Our findings of age-dependent cerebellar volume associations with cognitive scores in the amyloid-positive AD individuals who were younger than their older counterparts may reflect a survivor bias and lack of statistical power. Further, AD Aβ− patients may reflect a non-Alzheimer’s pathophysiologic process, such as suspected non-Alzheimer pathology (SNAP), characterized by heterogenous and potentially worse risk progression profiles (Wisse et al., 2018).

In spite of these limitations, we observed cerebellar volume-related mitigation of cognitive decline across healthy older adults and individuals along early stages of cognitive impairment and amyloid accumulation. These findings have profound implications for the identification of individuals at risk for AD dementia, given that the cerebellar protective effect may mask clinical symptoms in individuals whose robust cerebellar structures belie neocortical atrophy. We therefore urge researchers to consider cerebellar volume in their diagnostic criteria for neurodegenerative disease. In conclusion, our study highlights the cerebellum as a significant modulator of cognitive aging, especially in populations at greater genetic risk for Alzheimer’s disease. The cerebellum’s role in cognitive resilience, and its alignment with the threshold model of brain reserve, underscore its importance in understanding and potentially mitigating age-related cognitive decline.

## Data Availability

Deidentified participant data can be requested upon publication from the authors. Data sharing may necessitate appropriate data sharing agreements from the individual publicly accessible databases, including LONI for ADNI and HCP data (https://ida.loni.usc.edu/), and biobankUK for UK Biobank data (https://biobank.ctsu.ox.ac.uk).

## Author Contributions

FdU conceptualized and designed the study. FdU, JSeidlitz, JM, VZ, RRG, VW, RAIB, AFAB, and PV contributed to data access, curation, and processing. FdU, EF, JSeidlitz, SS-HW, PV, and JG contributed to data analysis. FdU, EF, JSeidlitz, JDC, JSepulcre, SS-HW, PV, and JG contributed to data interpretation and guided study progress. FdU, SSH-W, PV, and JG were responsible for writing the manuscript.

