## Supplemental Material for "A protective role for the cerebellum in cognitive aging"

*d'Oleire Uquillas et al., 2024*

**Table of Contents**

| Supplemental Methods | … | pgs. | 2 |
| --- | --- | --- | --- |
| Supplemental Figures | … | pgs. | 3 |

**Supplemental Methods**

**Montreal Cognitive Assessment**

The MoCA is a quick screening instrument for mild cognitive impairment, ranging from 1-30 points, and administered in 10-15 minutes, during which different cognitive domains are measured: Short-term memory recall, delayed memory recall, visuospatial abilities (clock drawing, 3D cube copy), processing speed (Trails Making B; Reitan, 1958), working memory (sustained attention task, serial subtraction task, and digits forward and backward memory task), language (3-item confrontation naming task, repetition of 2 syntactically complex sentences, and a verbal fluency task), and orientation (to time and place where being evaluated).

**Clinical Diagnosis and Grouping in ADNI**

For ADNI participants, a clinical diagnosis of MCI or AD was ascertained by a consensus of trained clinicians, considering cognitive scores including the MoCA, the Clinical Dementia Rating (CDR) scale (0 for normal cognitive function, 0·5 for mild cognitive impairment, 1 for mild dementia, 2 for moderate dementia, and 3 for severe and advanced dementia), and the Alzheimer’s Disease Inventory. Our ADNI analysis considered a total of five hundred and sixty-five CDR=0, six hundred and seventy-two CDR=0·5, one hundred forty-seven CDR=1, thirty-three CDR=2, and four CDR=3 individuals. The ADNI sample was stratified by ‘clinically normal’ (CN), mild cognitive impairment (MCI), and AD dementia clinical diagnosis. We opted to include individuals with a ‘subjective cognitive concerns’ in the CN group, given previous reports showing that optimal sub-group classification of ADNI data using spectral embedding, multidimensional scaling, UMAP, and t-SNE, is best captured by CN, MCI, and AD diagnostic groupings (van der Haar, et al., 2023).

**Supplemental Figures**

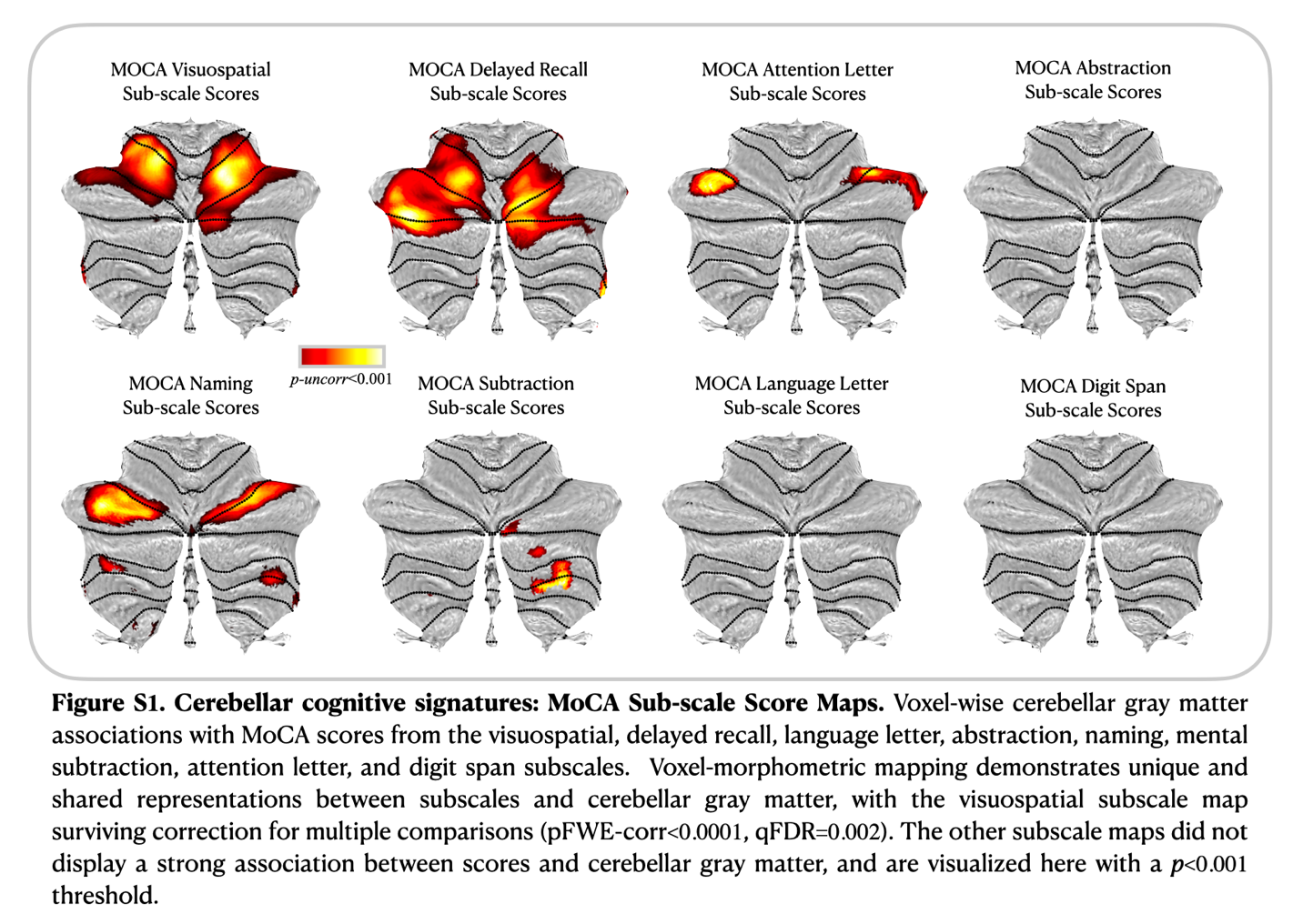

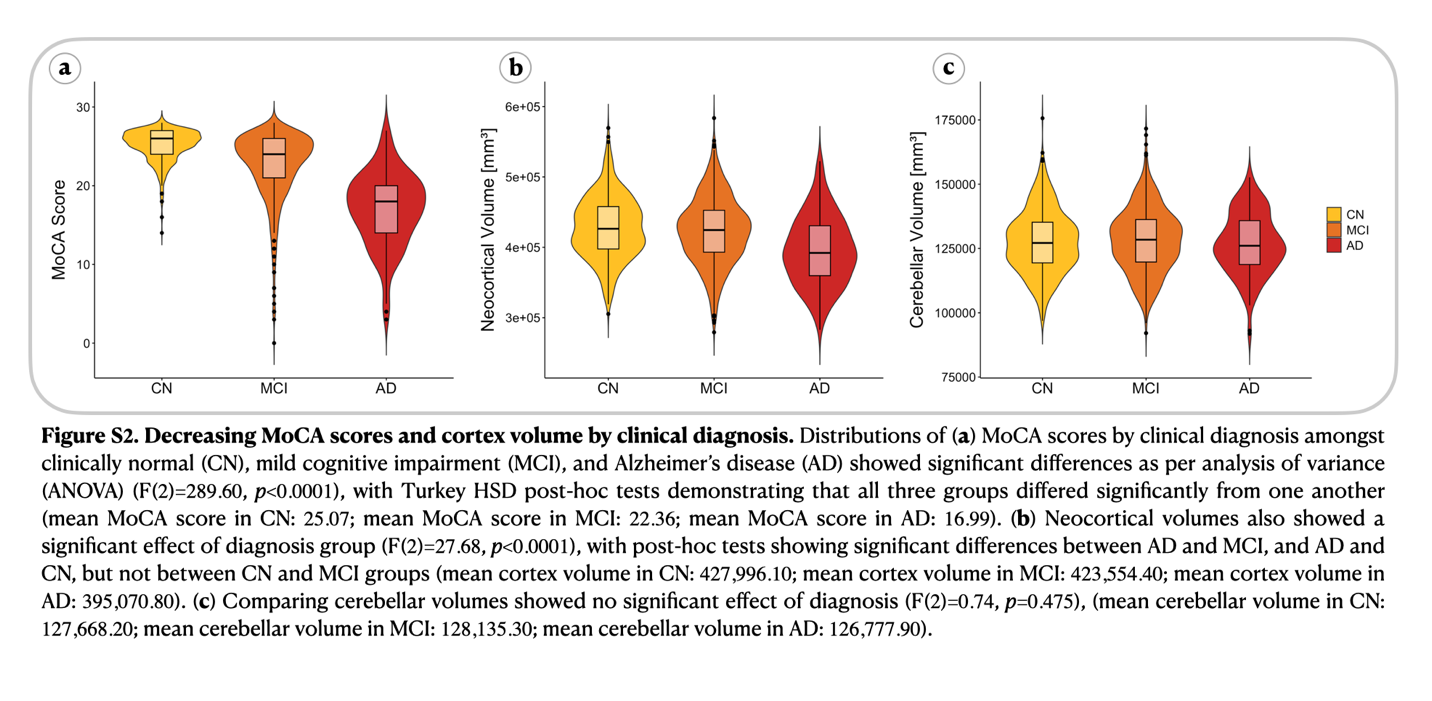
